# Paracrine regulations of IFN-γ secreting CD4^+^ T cells by lumican and biglycan are protective in allergic contact dermatitis

**DOI:** 10.1101/2024.10.20.619307

**Authors:** George Maiti, Jihane Frikeche, Cynthia Loomis, Shukti Chakravarti

**Author notes:** Corresponding authors: Shukti Chakravarti, PhD and George Maiti, PhD, **Email:**.

## Abstract

The extracellular matrix (ECM) is known to regulate innate immune cells but its role in T cell functions is poorly understood. Here, we show a protective role for ECM proteoglycans, lumican and biglycan in hapten-induced contact dermatitis that is achieved through limiting proinflammatory CD4^+^T cells. Lumican and biglycan-null mice develop significant inflammation with greater numbers of CD4^+^T cells in hapten-challenged ear pinnae, while their draining lymph nodes show increased T-bet-STAT1 signaling, Th1 commitment, and IFN-γ secreting CD4^+^T cell proliferation. Wild type mouse lymph node fibroblastic reticular cells secrete lumican, biglycan and decorin, a related proteoglycan, while none are expressed by naive or activated T cells. *In vitro*, lumican and biglycan co-localize with LFA-1 on T cell surfaces, and all three proteoglycans suppress LFA-1 mediated T cell activation. Overall, this study elucidates a novel paracrine regulation of Th1 cells by ECM proteoglycans to limit inflammation and tissue damage.

## Introduction

Among all types of contact dermatitis, allergic contact dermatitis (ACD) is a common worldwide problem with a prevalence of 20% in the general population, and about 16% in children (*1*). ACD develops from repeated exposure of the skin to environmental allergens called haptens, and involves a type IV delayed-type hypersensitivity response that derives from both the innate and adaptive immune system. Significant studies have focused on the immune regulatory signals that drive inflammation and dermal tissue damage, but few studies have examined the dermal extracellular matrix (ECM) as an active regulator of immune inflammatory responses. ACD develops in two phases: sensitization and elicitation. During the sensitization phase of repeated exposure to haptens, the skin goes through a pro-inflammatory and inflammatory phase. This initial pro-inflammatory milieu activates the epithelium, skin dermal dendritic cells (DCs) and epidermal Langerhans cells (LCs) that migrates toward the draining lymph node (dLN). Within the dLN, the DCs and LCs induces hapten-specific T cell activation and proliferation. The adaptive-immune response drives the elicitation phase where the hapten-specific activated CD4^+^ and CD8^+^ T cells migrate to the target tissues upon re-exposure to hapten causing the skin lesions. These infiltrating T cells secrete cytokines like IFN-γ along with other chemokines that induce migration of neutrophils to the skin aggravating tissue inflammation and damage (*2–4*). The ECM is an integral part of these early innate and late adaptive immune responses in the skin and the LN stroma that has not been explored well (*5, 6*). Therefore, we investigated functions of small leucine-rich repeat proteoglycans (SLRPs) of the ECM in ACD. We focused on the prototypical SLRPs, lumican, biglycan and decorin in ACD using a mouse model of contact hypersensitivity (CHS) (*2*), where mice were sensitized with the hapten, 2,4-Dinitrofluorobenzene (DNFB) followed by elicitation of the ear tissue with DNFB-challenge.

SLRPs contain highly interactive leucine-rich repeat (LRR) motifs in their core proteins that associate with fibrillar collagens in the ECM and regulate fibril assembly and connective tissue architecture and integrity (*7–10*). Lumican, biglycan and decorin, bind fibrillar collagens (*11*), and gene-targeted null mice lacking these proteoglycans harbor defective collagen fibril architecture in the skin, tendon, cartilage, cornea, and vascular walls with associated functional loss in these tissues (*12–15*). While these early studies established SLRPs as structural collagen regulating components of the ECM, later studies began to show that many SLRPs are induced during injury, infection, and inflammation and secreted by activated fibroblasts (*16–18*). These *de novo* synthesized proteoglycans and core proteins, as well as those released from an injury-responsive remodeling ECM appear to interact with immune cells to shape disease. For instance, lumican binds the co-receptor CD14 and to Cav1 in lipid rafts, to enhance TLR4 stability and TLR4 mediated pro-inflammatory signaling in macrophages and DCs (*18–20*). Biglycan, reportedly promotes TLR4 signals as a ligand itself, and both null strains (*Lum*^-/-^, *Bgn^-/-^* and *Bgn^-/^*°) are hyporesponsive to bacterial lipopolysaccharide-mediated sepsis (*19, 21*). We also found that lumican and biglycan restrict TLR9 response. Both bind CpG DNA, and lumican colocalizes with CpG DNA in TLR9-poor early endosomes, suggesting their regulation of TLR9 signals may in part be through ligand sequestration (*18, 22*). Suppression of TLR9 response may be an important means of limiting autoimmune-leaning response to self-DNA in CHS as well. While here we focus on the cross talk between SLRPs and T cells, their role in early innate immune inflammatory signals help to regulate the pro-inflammatory milieu in dermal tissues in the CHS mouse model of ACD.

All three SLRPs have been implicated in inflammatory diseases, like sepsis and keratitis, and autoimmune diseases, such as multiple sclerosis and nephritis (*16, 18, 20, 23*). In addition to being major components of dermal connective tissues, we show their specific locations in secondary lymphoid tissues. Several SLRP genes are expressed by fibroblastic reticular cells (FRCs) of the mouse LN, spleen and Peyer’s patches (*5, 24–26*). FRCs secrete many factors that are critical in regulating the LN microenvironment and T cell functions. For example, the chemoattractant CCL19 and CCL21, secreted by FRCs regulate naïve T cell trafficking. They also regulate T cell homeostasis and priming by facilitating T-DC interactions (*27, 28*). In mice, FRCs suppress T cell activation and proliferation by producing prostaglandin E2 (PGE2) and upregulating nitric oxide production, respectively (*29, 30*). In humans, LN-derived FRCs also suppress T cell activation and differentiation (*31*). Thus far, no studies have investigated whether SLRPs produced by FRCs regulate T cell functions.

Here we show that lumican- (*Lum^-/-^*) and biglycan- (*Bgn^-/-^* and *Bgn^-/0^*) knockout mice experience heightened ear inflammation in the CHS mouse model. The draining lymph node (dLN) of *Lum^-/-^*and *Bgn^KO^* mice show significantly higher numbers of proliferating CD4^+^ T cells, which are mostly IFN-γ secreting Th1 cells. Using an OT-II adoptive transfer model, we found that lumican, biglycan, and decorin suppress the activation of adoptively transferred cells in the dLN of wild type mice. Lumican, biglycan, and decorin secreted by the FRCs interact with the β2 integrin subunit of LFA-1 and compete with ICAM-1 to suppress the CD4^+^ T cell activation. Our findings underscore a paracrine role for SLRPs secreted by FRCs in homeostasis and inflammation, in CD4^+^ T cell regulations, and as promising therapeutic proteins or peptides to restrict Th1-driven tissue damage.

## Results

### Increased influx of CD4^+^ T cells and ear inflammation in hapten-challenged *Lum^-/-^*and *Bgn^KO^* mice

Using wild type (WT) and null mice systemically lacking lumican or biglycan, we tested their role in the CHS model which involves a classic Th1-driven skin allergic reaction (*2*). *Lum^-/-^*, *Bgn^-/-^* and *Bgn^-/0^* (referred to as *Bgn^KO^,* hereafter), and WT mice were sensitized by painting their belly with the hapten, 2, 4-dinitrofluorobenzene (DNFB) on days 0 and 1, then challenged on day 5 by exposing their ear pinnae to DNFB **(Fig. 1A)**. Ear swelling was measured before the challenge on day 5 and days 6 and 7 using a digital vernier caliper. All unchallenged mice had comparable ear thickness on days 5, 6 and 7. However, challenged mice on days 6 and 7 of the timeline, show thickened ear pinnae, and this is significantly (*p* < 0.01) higher in *Lum^-/-^* and *Bgn^KO^* mice (0.55 ± 0.014 and 0.56 ± 0.022 mm, respectively) compared to WTs (0.37 ± 0.013 mm) **(Fig. 1B)**. H and E staining of ear pinnae sections of unchallenged mice of all three genotypes show comparable tissue architecture and thickness, whereas after the challenge *Lum^-/-^* and *Bgn^KO^* show increased edema and immune cell infiltrates compared to WT (**Fig. 1C**). WT mice show increased immunostaining of lumican in the dermal layers, while biglycan is markedly increased in the epithelium as well **(Fig. S1A and S1B)**. Using multiplexed fluorescent immunohistochemistry, we next analyzed the type of immune cells that infiltrate the ear tissue of challenged mice at day 7. We detected neutrophils and macrophages by immunostaining for Ly6G and F4/80, respectively.

**Fig. 1.**
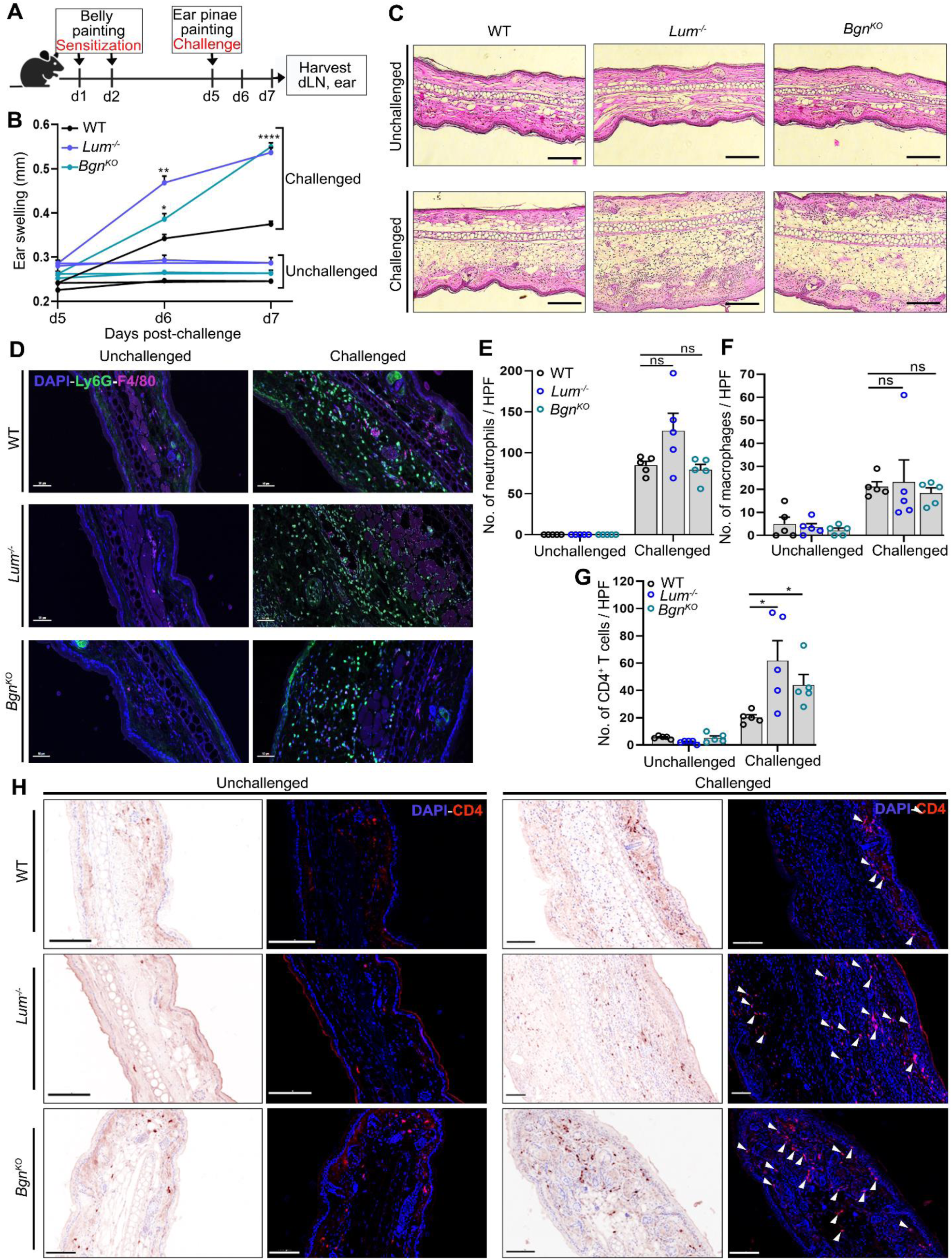
Lumican and biglycan suppress ear swelling in CHS. (**A**) Schematic showing the CHS model. (**B**) Quantification of ear swelling (n = 9 – 13 mice) from 2 independent experiments. (**C**) Representative histology images showing hematoxylin and eosin (H&E) staining of ear tissue sections from unchallenged and challenged WT, *Lum^-/-^* and *Bgn^KO^* mice. (**D**) Representative confocal images of ear tissue sections showing DAPI, Ly6G and F4/80 staining. (**E** and **F**) Cumulative data showing number of Ly6G^+^ neutrophils and F4/80^+^ macrophages in 5 high power field (HPF) images in (D) (n = 2 mice/genotype) from two independent experiments. (**G**) Representative confocal images of ear tissue sections showing nuclear stain, DAPI and CD4 staining. The white arrows indicate the infiltrating CD4^+^ T cells. The corresponding bright-field images show the chromogenic HRP/DAB staining specific for CD4. (**H**) Cumulative data showing the number of CD4^+^T cells in 5 HPF images in (D) (n = 2 mice/genotype) from two independent experiments. Scale bars, 50 µm (C) and 100 µm (D and H). The error bars represent mean ± SEM, two-way ANOVA followed by Sidak’s multiple comparisons test (B), and unpaired t test with Welch correction (E, F and G). *\*P < 0.05, **P < 0.01, ****P < 0.0001*, *ns, not significant.* See also figures S1, S2 and S3.

In the challenged group, the number of Ly6G^+^ neutrophils and F4/80^+^ macrophages were similar in the three genotypes **(Fig. 1D – 1F and S2A)**. Immunohistology show little ingress of CD4^+^ T cells in the ear pinnae dermis of all unchallenged mice, as expected. By contrast, all challenged mice showed increased CD4^+^ T cells, with visibly more staining of CD4^+^ T cells in sections of *Lum^-/-^* and *Bgn^KO^*ear pinnae (**Fig. 1G**). We next counted the numbers of CD4^+^ T cells from 5 high power field (HPF) images from two experiments (**Fig. 1H**). Challenged *Lum^-/-^* and *Bgn^KO^* mouse ear pinnae showed significantly higher numbers of CD4^+^ T cells compared to WTs. Foxp3^+^ CD4^+^ Tregs are known to suppress ear swelling in the CHS model (*32*). Therefore, we quantified infiltrating Foxp3^+^CD4^+^ Tregs and found a subset of CD4^+^ T cells to be Foxp3^+^, but overall, the numbers were comparable in WT, *Lum^-/-^* and *Bgn^KO^* challenged mice **(Fig. S2B and S2C)**. Altogether, CHS model shows greater ear inflammation and increased CD4^+^ T cells in the *Lum^-/-^* and *Bgn^KO^* mice compared to WTs.

### Increased proliferating CD4^+^ T cells in the draining lymph nodes of hapten-challenged *Lum^-^****^/-^* and *Bgn^KO^* mice**

Although we found higher numbers of CD4^+^ T cells in the *Lum^-/-^* and *Bgn^KO^* mouse ear pinnae, percentage of proliferating CD4^+^ T cells were similar in all three genotypes (**Fig. S3A and S3B)** Therefore, we wondered whether increased proliferation of CD4^+^ T cells in the ear dLN was contributing to the increase in CD4^+^ T cell seen in the inflamed ear tissues. We used flow cytometry to quantify the number of proliferating CD4^+^ and CD8^+^ T cells in the dLN of unchallenged and challenged mice on day 7 of the model. Total numbers of both CD4^+^ and CD8^+^ T cells were significantly higher in the dLN of challenged null mouse strains compared to WTs **(Fig. 2A and 2B)**, while percent of these cell types over total cells per dLN were similar **(Fig. S4A and S4B)**. Importantly, proliferating PCNA^+^CD4^+^ T cells were increased significantly in the dLN of *Lum^-/-^* and *Bgn^KO^*mice compared to WT **(Fig. 2C – 2E)**. However, proliferating PCNA^+^CD8^+^ T cells were similar in the dLN of challenged WT, *Lum^-/-^*and *Bgn^KO^* mice **(Fig. 2F and 2G)**. Altogether, our data demonstrates that the absence of Lum and Bgn in the dLN lead to increased CD4^+^ T cell proliferation in dLN in *Lum^-/-^*and *Bgn^KO^* mice, respectively.

**Fig. 2.**
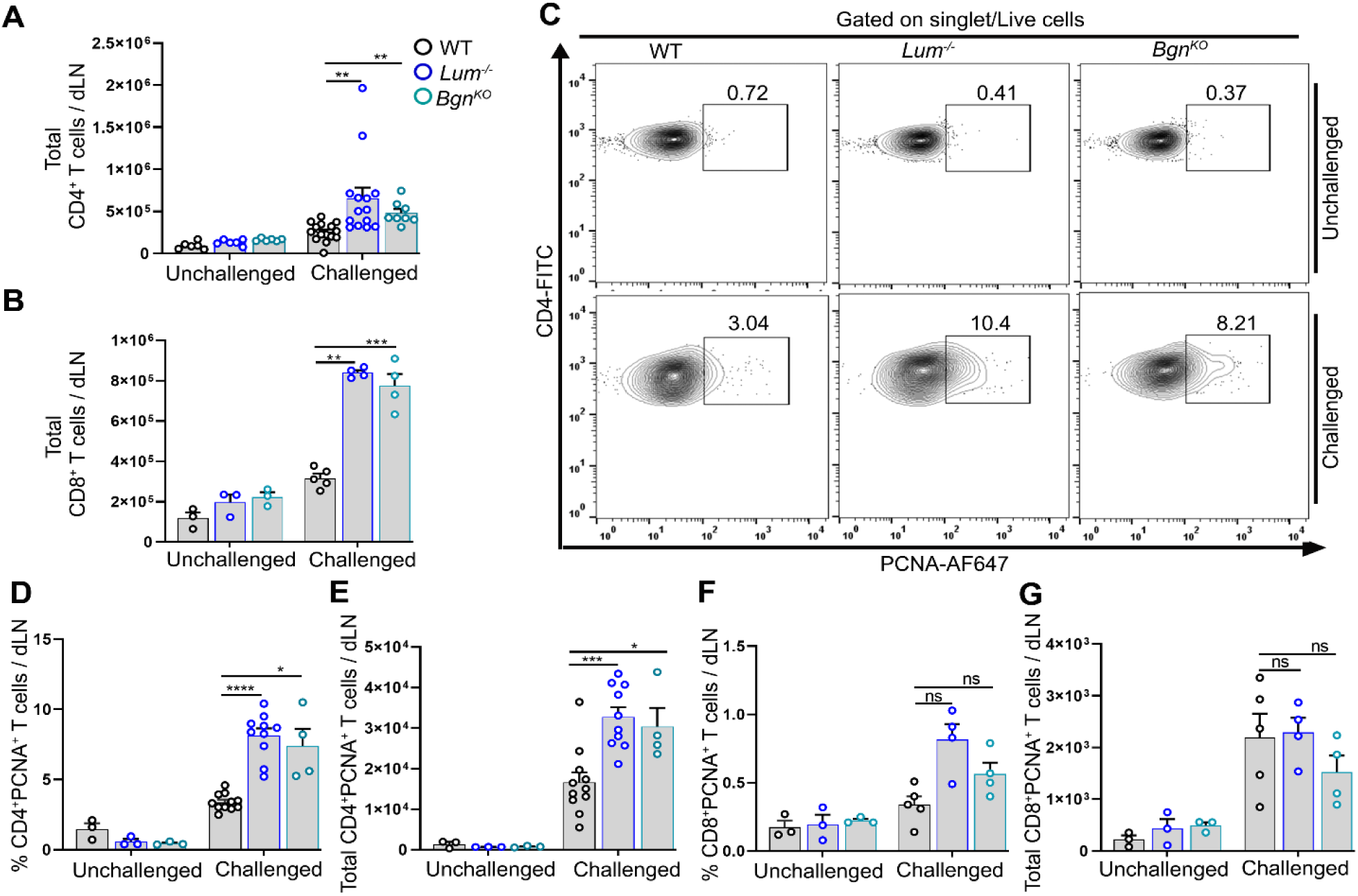
Increased CD4^+^ T cell proliferation in the draining LN of *Lum^-/-^* and *Bgn^KO^* mice. (**A** and **B**) The cumulative data shows the total number of CD4^+^ T cells (A) and CD8^+^ T cells (B) in the dLN of unchallenged and challenged WT, *Lum^-/-^* and *Bgn^KO^* mice (n = 6 – 14) from 3 independent experiments. (**C**) One representative contour plot from 2 independent experiments shows the frequency of proliferating PCNA^+^CD4^+^ T cells in the dLN. (**D** and **E**) The cumulative data show the percentage (D) and total number (E) of PCNA^+^CD4^+^ T cells in the dLN of unchallenged (n = 3) and challenged (n = 4 – 15) WT, *Lum^-/-^*and *Bgn^KO^* mice. (**F** and **G**) The cumulative data show the percentage (F) and total number (G) of PCNA^+^CD8^+^ T cells in the dLN (n = 3 -5 mice/genotype from 2 independent experiments). The error bars represent mean ± SEM, unpaired t test with Welch correction. *\*P < 0.05, **P < 0.01, ***P < 0.001, **** P < 0.0001, ns, not significant*. See also figure S4.

### Increased IFN-γ secreting CD4^+^ T Th1 cells and not Th17 cells in the draining lymph nodes of hapten-challenged *Lum^-/-^* and *Bgn^KO^* mice

Previous studies have shown that DNFB mediated CHS is primarily driven by IFN-γ producing CD4^+^ T (Th1) cells (*33, 34*). Therefore, we tested if the increased ear swelling in the *Lum^-/-^* and *Bgn^KO^*mice is due to increased Th1 or Th17 cells in the dLN. Cells extracted from the dLN of challenged mice were restimulated with PMA and ionomycin, and analyzed for Th1, Th2 and Th17 subsets by flow cytometry. We found significantly higher percentage of IFN-γ secreting CD4^+^ T cells in the dLN of *Lum^-/-^*(15.31 ± 1.39%) and *Bgn^KO^* (15.84 ± 1.0%) mice compared to WT (6.9 ± 0.45%) **(Fig. 3A and 3B)**. Moreover, the total number of (IFNγ^+^CD4^+^) Th1 cells is significantly increased in both null mouse strains compared to WT, suggesting a Th1 differentiation-limiting function for lumican and biglycan **(Fig. 3C)**. By contrast, IL-17 secreting CD4^+^ T (Th17) cells or IL-4 secreting CD4^+^ T (Th2) cells are similar in all three genotypes **(Fig. 3D – 3F)**.

**Fig. 3.**
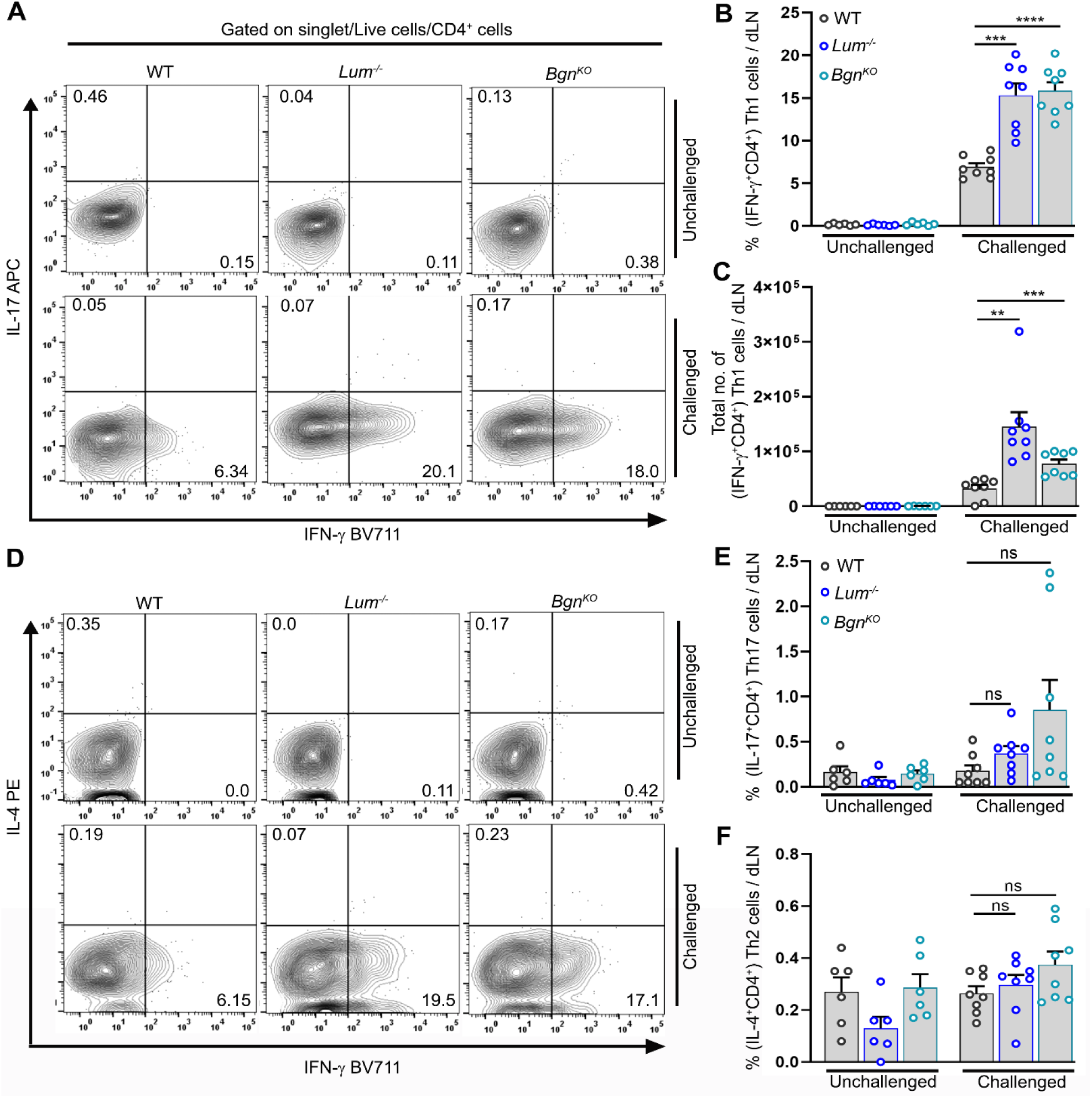
Elevated Th1 cells in the draining LN. (**A**) Representative contour plots showing the frequency of IFN-γ secreting CD4^+^ T (Th1) and IL-17 secreting CD4^+^ T (Th17) cells in the dLN on day 7 of the DNFB model. (**B** and **C**) The cumulative data shows the percentage (B) and total number (C) of Th1 cells in the dLN of unchallenged and challenged WT, *Lum^-/-^* and *Bgn^KO^* mice (n = 6 – 8 mice/genotype) from 2 independent experiments. (**D**) Representative contour plots showing the frequency of IL-4 secreting CD4^+^ T (Th2) cells in the dLN. (**E** and **F**) The cumulative data shows the percentage of Th17 (E) and Th2 (F) cells in the dLN (n = 6 – 8 mice/genotype from 2 independent experiments). The error bars represent mean ± SEM, unpaired t test with Welch correction. *\*\*P < 0.01, ***P < 0.001, **** P < 0.0001, ns, not significant*.

The transcription factor T-bet is critical for Th1 lineage commitment and is upregulated by IFN-γ (*33, 34*). Therefore, we checked the T-bet expression in CD4^+^ T cells in the dLN of unchallenged and challenged *Lum^-/-^*, *Bgn^KO^* and WT mice by flow cytometry **(Fig. 4A)**. Our data shows that in the challenged group, the percentage of T-bet expressing CD4^+^ T cells is significantly higher in the *Lum^-/-^* (3.42 ± 0.36%) and *Bgn^KO^*(5.99 ± 1.01%) compared to the WT (2.64 ± 0.23%) mice **(Fig. 4B)**. Similarly, the total number of T-bet^+^CD4^+^ T cells in the dLN of *Lum^-/-^*and *Bgn^KO^* mice are significantly higher than WTs **(Fig. 4C)**. The numbers of T-bet^+^ CD4^+^ T cells in unchallenged mice of all three genotypes are low and comparable. We further analyzed the IFN-γ-STAT1-T-bet signaling axis by quantifying phosphorylated STAT1 (pSTAT1) levels in the dLN by immunoblotting. The phospho-STAT1 band was stronger in the dLN of *Lum^-/-^*and *Bgn^KO^* mice, consistent with a significant increase in IFN-γ signaling in the null strains compared to WT **(Fig. 4D, 4E and S5A – S5B)**. Taken together, these observations indicate that lumican and biglycan suppress Th1 differentiation and proliferation by inhibiting the IFN-γ-STAT1-T-bet signaling axis.

**Fig. 4.**
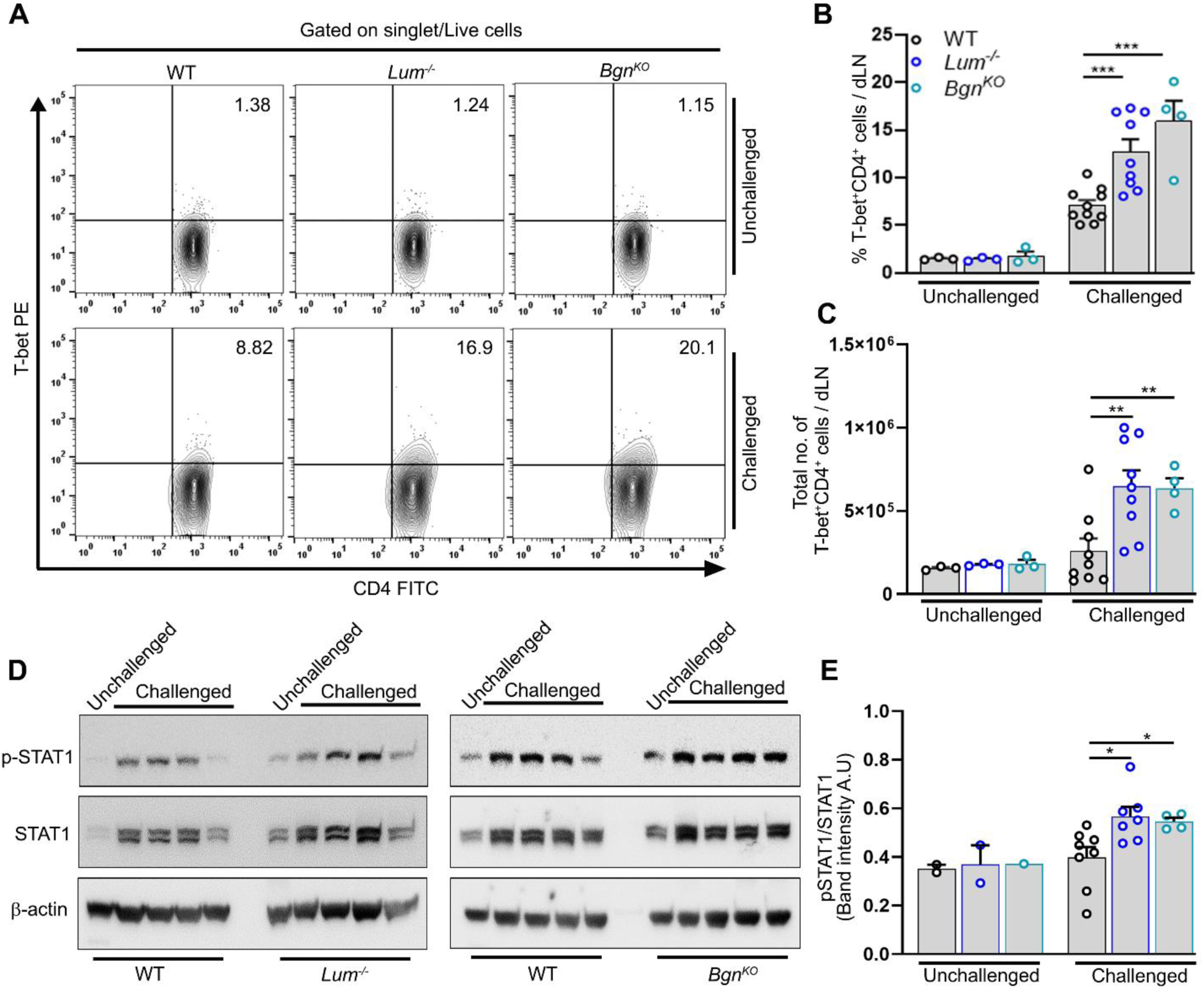
Increased T-bet and phospho-STAT1 levels in Th1 cells in *Lum^-/-^* and *Bgn^KO^* mice. (**A**) Representative contour plots showing the frequency of T-bet^+^CD4^+^ T (Th1) cells in the dLN. (**B** and **C**) The cumulative data shows the percentage (B) and total number (C) of T-bet expressing Th1 cells in the dLN of unchallenged and challenged WT, *Lum^-/-^* and *Bgn^KO^* mice (n = 3 – 10 mice/genotype) from 2 independent experiments. (**D**) Representative immunoblot of dLN lysates from unchallenged and challenged WT, *Lum^-/-^* and *Bgn^KO^* mice. (**E**) The cumulative data shows phospho-STAT1 level normalized to STAT-1 level from 2 independent experiments (n = 1 – 8 mice/genotype). The error bars represent mean ± SEM, unpaired t test with Welch correction. *\*P < 0.05, **P < 0.01, ***P < 0.001*. See also figure S5.

### Increased activation of adoptively transferred OT-II CD4^+^ T cell in OVA-challenged *Lum^-/-^* **and *Bgn^KO^* mice**

Given that DNFB challenged *Lum*^-/-^ or *Bgn*^KO^ mouse LNs show increased proliferating CD4^+^ T cells, we wondered if SLRPs actually limit the activation of CD4^+^ T cells. We used an *in vivo* adoptive transfer model to investigate how these SLRPs directly impact antigen specific CD4^+^ T cell activation (*35*). GFP^+^ OT-II TCR transgenic CD4^+^ T cells (10^6^ cells) were adoptively transferred to *Lum^-/-^*, *Bgn^KO^* and WT recipient mice. The OT-II cells recognize ovalbumin (OVA) peptide OVA^323-339^ presented by the MHC-II molecule on APCs (*36, 37*). After 24 h, one hind foot pad was injected with OVA emulsified in incomplete Freund’s adjuvant (IFA), and the other foot pad received the control PBS injection.

On day 2, the footpad draining popliteal LNs were isolated to quantify the number of activated OT-II cells **(Fig. 5A)**. The number of OT-II cells in the inflamed dLN was significantly higher in *Lum^-/-^* and *Bgn^KO^* compared to WT **(Fig. 5B and 5C)**. However, the percent OT-II cells were comparable among the three genotypes **(Fig.S6A)**. Next, we assayed CD69 expression in CD4^+^ OT-II cells to determine their activation status. Absolute numbers and percentage of CD69^+^CD4^+^OT-II cells were considerably higher in *Lum^-/-^* and *Bgn^KO^* mice than WT, indicating increased activation in the absence of lumican or biglycan **(Fig. 5D – 5F)**.

**Fig. 5.**
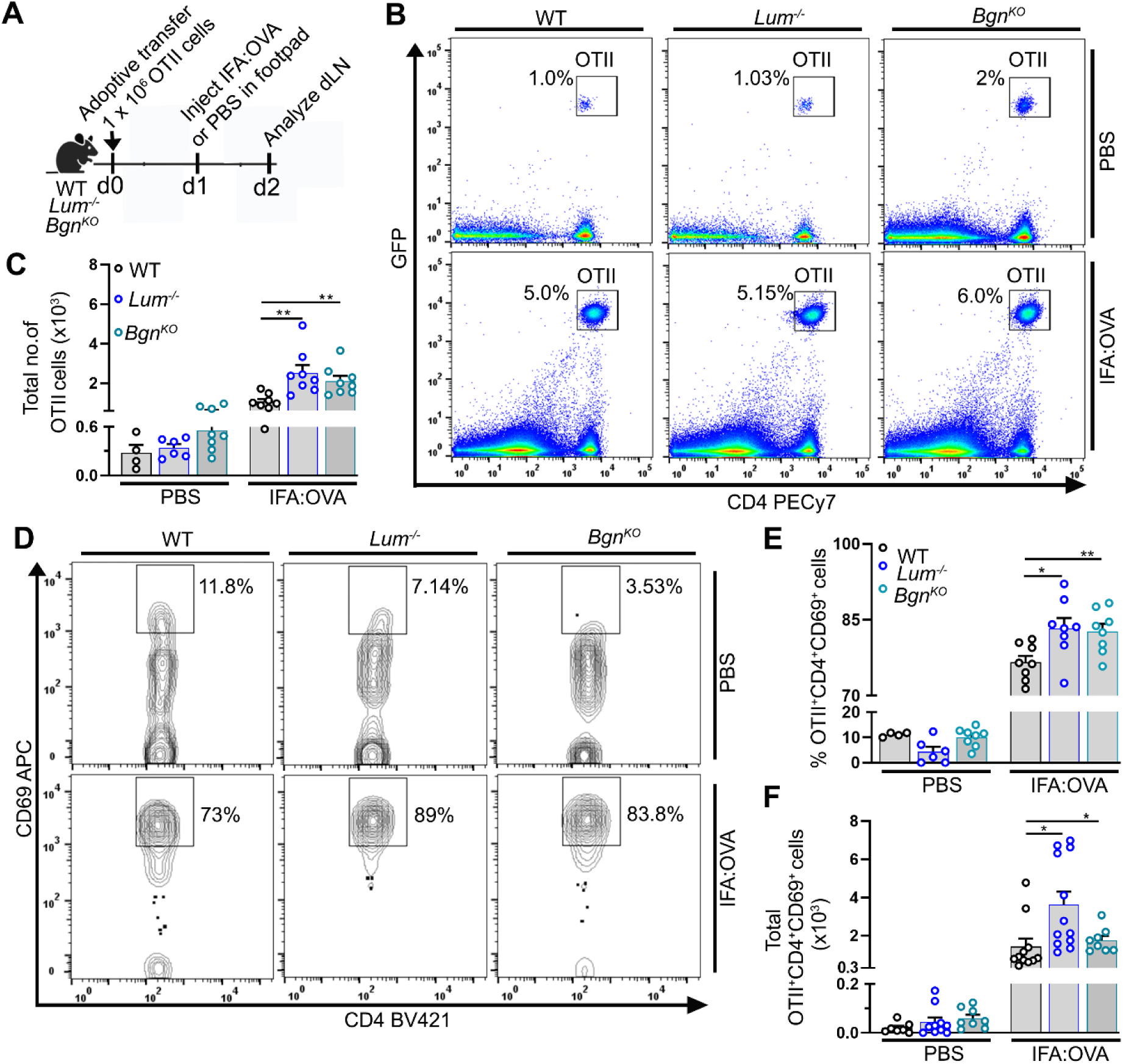
Lumican and biglycan suppress CD4^+^ T cell activation. (**A**) Assay for *in vivo* activated (Act) CD4^+^ T cells using the OT-II adoptive transfer model. (**B**) Representative flow cytometry dot plots showing OT-II (GFP^+^CD4^+^) cells in the dLN after IFA-OVA or PBS injection in WT, *Lum^-/-^* and *Bgn^KO^* mice. (**C**) Cumulative data shows total number of transferred OT-II cells in the dLN (n = 4 – 8 mice/genotype) from 2 independent experiments. (**D**) Representative contour plots showing percentage of activated OT-II cells (CD69^+^CD4^+^ T cells) in the dLN. Gated on singlet and live OT-II cells. (**E** and **F**) Cumulative data shows the percentage (E) and total number (F) of activated OT-II cells (CD69^+^OT-II^+^CD4^+^ T cells) in the dLN after injected with either IFA-OVA or PBS in WT, *Lum^-/-^* and *Bgn^KO^* mice (n = 4 – 12 mice/genotype) from atleast 2 experiments. The error bars represent mean ± SEM, unpaired t test with Welch correction. *\*P < 0.05, **P < 0.01*. See also figures S6 and S7.

Decorin is another related SLRP that is abundantly present in the ECM and binds to TLR-2 and -4 on macrophages to promote the production of pro-inflammatory factors (*23*). Decorin null mice compared to WT mice were less hyperresponsive with lower inflammation and airway remodeling in an OVA-challenged allergen-induced asthma model (*38*). Therefore, we tested the role of decorin in regulating CD4^+^ T cell activation using the OT-II model in *Dcn^-/-^* mice **(Fig. S6B)**. Although dLN size and total cells **(Fig. S6C and S6D)** and the percentage and number of OT-II cells **(Fig. S6E – S6G)** were comparable in *Dcn^-/-^*and WT mice, the percentage and number of activated CD69^+^ CD4^+^ OT-II cells were significantly higher in the *Dcn^-/-^* mice, as we noted in the *Lum^-/-^* and *Bgn^KO^*mice **(Fig. S6H – S6J).**

### Exogenous lumican and biglycan suppress CD4^+^ T cell activation *in vitro* via LFA-1

Entry of naïve T cells into the dLN depends on the interaction between LFA-1 on the T cell membrane and its ligand ICAM-1 on the endothelial cells. LFA-1, a heterodimeric (*α*L/*β*2 or CD11a/CD18) integrin, actively regulates the interactions between CD4^+^ T and DC at the immunological synapse (IS) (*39, 40*). Formation of a stable IS induces adequate engagement for TCR with antigen-MHC complex that initiates the phosphorylation of Lck and ZAP-70, and subsequently leads to full T cell activation These events culminate in the activation, proliferation, and differentiation of naïve T cells (*41, 42*). Therefore, we next sought to test if recombinant (r) lumican (rLum) and biglycan (rBgn) suppress the TCR downstream signaling strength by quantifying the phosphorylation level of Lck and ZAP-70. The phosphorylated ZAP-70 level remained significantly low in activated CD4^+^ T cells pretreated with rLum or rBgn. CD4^+^ T cell suppression can be detected as early as 15 min of activation **(Fig. 6A)**. However, the phosphorylated Lck levels were comparable at early time points but significantly reduced at 60 min of activation in rLum or rBgn pretreated CD4^+^ T cells **(Fig. 6B)**. In a transwell migration assay, we further tested the recombinant proteoglycans ability to suppress chemotaxis of naïve CD4^+^ T cells toward recombinant CXCL19. Our data shows that all three SLRPs significantly inhibit the chemotaxis of CD4^+^ T cells compared to PBS treated controls **(Fig. S7A and S7B)**. Overall, these results indicate that lumican, biglycan and decorin can restrict CD4^+^ T cell activation and chemotaxis in the dLN.

**Fig. 6:**
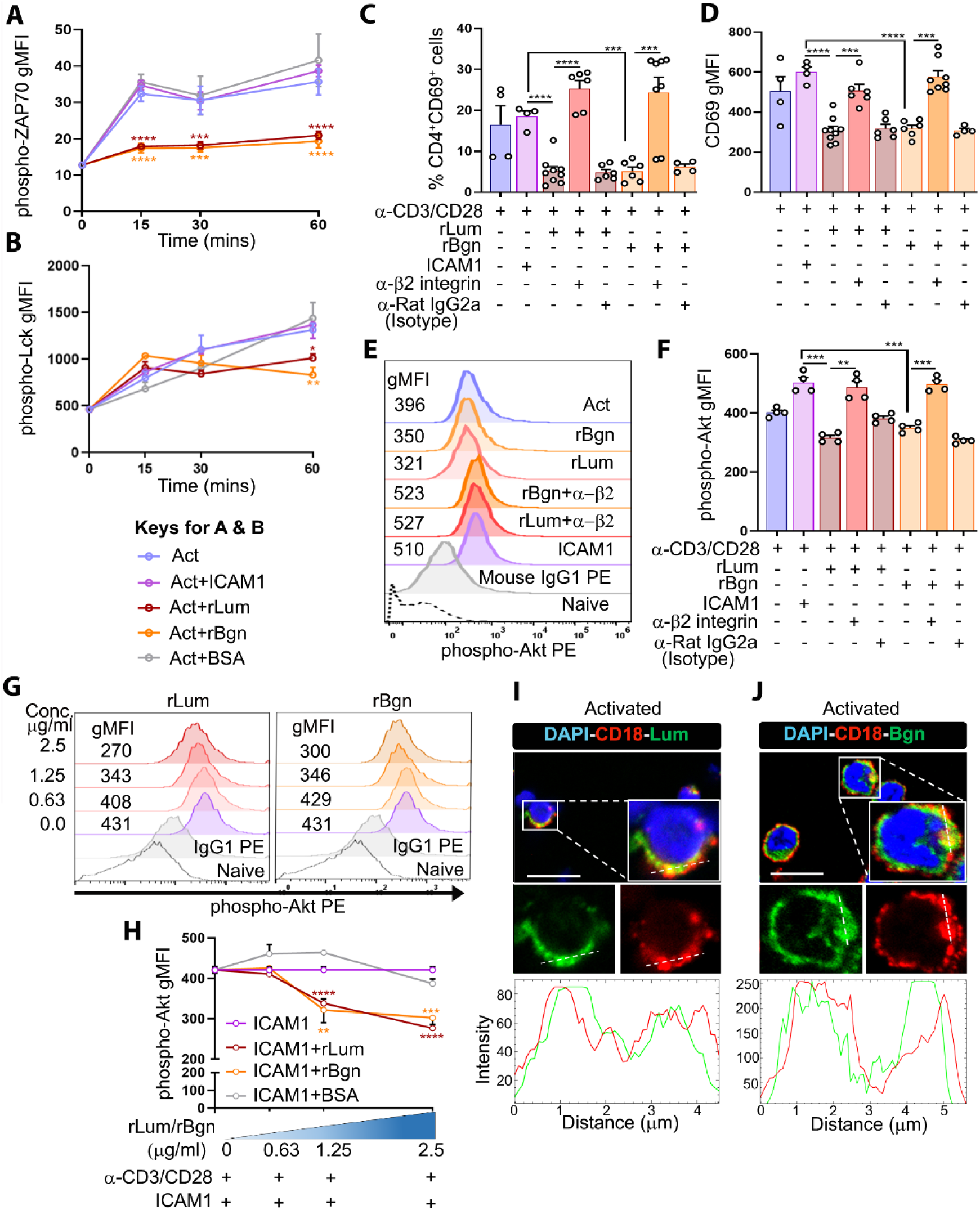
Lumican and biglycan colocalize with LFA-1 and suppress CD4^+^ T cell functions. (**A** and **B**) Cumulative data from 2 independent experiments show geometric mean fluorescence intensity (gMFI) of phospho-ZAP-70 (A) and phospho-Lck (B) levels in activated CD4^+^ T cells treated with ICAM-1 or rLum or rBgn or BSA. (**C** and **D**) Cumulative graph from 2 independent experiments shows the percentage (C) of activated CD69^+^CD4^+^ T cells and surface expression (gMFI) of CD69 (D). (**E**) Representative histograms show the phospho-Akt-1 level after 30 min in activated CD4^+^ T cells with various treatments as indicated in the plot. (**F**) Cumulative gMFI from 2 independent experiments shows the phospho-Akt-1 levels in activated CD4^+^ T cells treated with either rLum or rBgn or ICAM-1 alone or in combination with anti-β2 integrin antibody. Anti-Rat IgG2a is used as an isotype control. (**G**) Representative histograms show the phospho-Akt-1 level in activated CD4^+^ T cells with increasing concentration of soluble rLum or rBgn or BSA. (**H**) Cumulative gMFI from 2 independent experiments shows the phospho-Akt-1 levels in activated CD4^+^ T cells as a measure of competition between rLum, or rBgn and ICAM-1 for LFA-1 binding. (**I** and **J**) Representative confocal image from 2 independent experiments shows colocalization of CD18 with rLum (I) and rBgn (J). The histogram depicts cross-line scans of the fluorescence intensities of the merged panels. Scale bars, 100 µm. Each dot represents technical replicates (C, D and F). The error bars represent mean ± SEM, unpaired t test with Welch correction. *\*P < 0.05, **P < 0.01, ***P < 0.001, ****P < 0.0001*. See also figure S8.

Our earlier biochemical study showed that lumican binds to the β2 integrin subunit on polymorphonuclear neutrophils (*43*). The β2 (CD18) and αL (CD11a) subunit heterodimerize to form the LFA-1 receptor complex, which is robustly present on T cell membranes and is critical for T cell activation, proliferation, and Th1 differentiation (*44, 45*). Therefore, we asked if lumican and biglycan restrict CD4^+^ T cell activation by interacting with the β2 subunit of LFA-1. To investigate this, we tested suppression of CD4^+^ T cell activation (CD69) by the recombinant proteoglycans *in vitro* in the presence of anti-β2 integrin blocking antibody. Indeed, pretreatment of CD4^+^ T cells with anti-β2 blocking antibody could rescue the CD69 level compared to either rLum or rBgn treated cells **(Fig. 6C and 6D)**. Similarly, CD4^+^ T cells pretreated with anti-β2 integrin antibody rescued the rDcn mediated suppression of activation **(Fig. S8A and S8B)**. In fact, the surface CD69 levels was similar to that attained by treatment with ICAM-1, the natural ligand for LFA-1. CD4^+^ T cells pretreated with isotype IgG (anti-Rat IgG2a) as a control could not rescue the suppressive effect of lumican, biglycan, and decorin. These observations suggest that three proteoglycans mediate CD4^+^ T regulations through LFA-1.

Earlier studies show that binding of LFA-1 to ICAM-1 leads to the intracellular phosphorylation and activation of Akt-1 (*46*). Therefore, using *in vitro* activated CD4^+^ T cells we quantified intracellular phospho-Akt-1 in the presence of ICAM-1 and recombinant SLRPs to test their effects on this axis. All three proteoglycans reduced phospho-Akt-1 in a dose dependent manner at concentrations between 1.25 and 2.5 µg/ml. Here bovine serum albumin (BSA) as a non-SLRP control protein did not affect phospho-Akt-1 levels **(Fig. 6E – 6H and S8C – S8F)**.

This suggests that by binding to LFA-1 these proteoglycans are likely interfering with LFA-1-ICAM-1 interactions and subsequent phosphorylation of Akt-1. We next used immunohistology and confocal microscopy to tested colocalization of lumican and biglycan with the β2 subunit (CD18) on naïve and activated CD4^+^ T cells. Indeed, we detected strong colocalization of lumican and biglycan with CD18 on the activated CD4^+^ T cell membrane, which was not detectable on naïve CD4^+^ T cells **(Fig. 6I – 6J and S8G – S8H)**. Altogether, our results suggest that lumican and biglycan interact with the β2 subunit of LFA-1, and thereby restrict CD4^+^ T cell activation.

### Lumican, biglycan, and decorin secreted by the fibroblastic reticular cells in lymph node niche are likely regulators of CD4^+^ T cells

Previous transcriptomic studies show that LN stromal cells express lumican, biglycan, and decorin genes (*5, 24*). We localized lumican, biglycan, and decorin in different LN compartments by immunohistochemistry. Immunostaining of LN from WT mice shows that lumican and biglycan are extensively present throughout the capsule (C), sub-capsular sinus (SC), inter-follicular region (IFR) and T cell rich para-cortex region (T) and to a lesser extent in the B cell follicle (B) **(Fig. 7A – 7C and S9A – S9B)**. We further determined if lumican and biglycan colocalize with a stromal fibroblast marker, podoplanin (PDPN), and the T cell marker, CD3. Biglycan shows robust colocalization with PDPN, whereas, lumican shows weaker colocalization with PDPN^+^ fibroblastic reticular cells (FRCs). Both colocalize with CD3 to a lesser extent than PDPN, while neither colocalize with B220^+^ B cells **(Fig. 7D)**. Lumican and biglycan are also present in the spleen, the largest secondary lymphoid organ, and highly enriched in the capsule, trabeculae, red pulp and the splenic sinusoids **(Fig. S9C)**. Used as negative controls, *Lum^-/-^*and *Bgn^KO^* mouse LN sections showed little to no immunoreactivity **(Fig. 7E and 7F)**. Immunostaining for decorin in wild type mouse LNs localizes it in the capsule and subcapsular sinus connective tissues, with lesser staining of the IFR and T cell zone, and *Dcn^-/-^* mice LN, used as negative controls, show no immunoreactivity **(Fig. S10A-S10B)**. Decorin shows the strongest colocalization with PDPN^+^ FRCs, as corroborated by Pearson’s coefficient analysis between PDPN and decorin (r = 0.8), or lumican (r = 0.52) or biglycan (r = 0.6) **(Fig. S10C)**. In the spleen, decorin is predominantly enriched in the capsule, trabeculae, red pulp, and splenic sinusoids **(Fig. S10D)**. Taken together, these observations suggest that lumican, biglycan, and decorin are secreted by PDPN^+^ FRCs to various extents, and localize within distinct niches in the LN and spleen where they may regulate niche-specific immune cell functions.

**Fig. 7:**
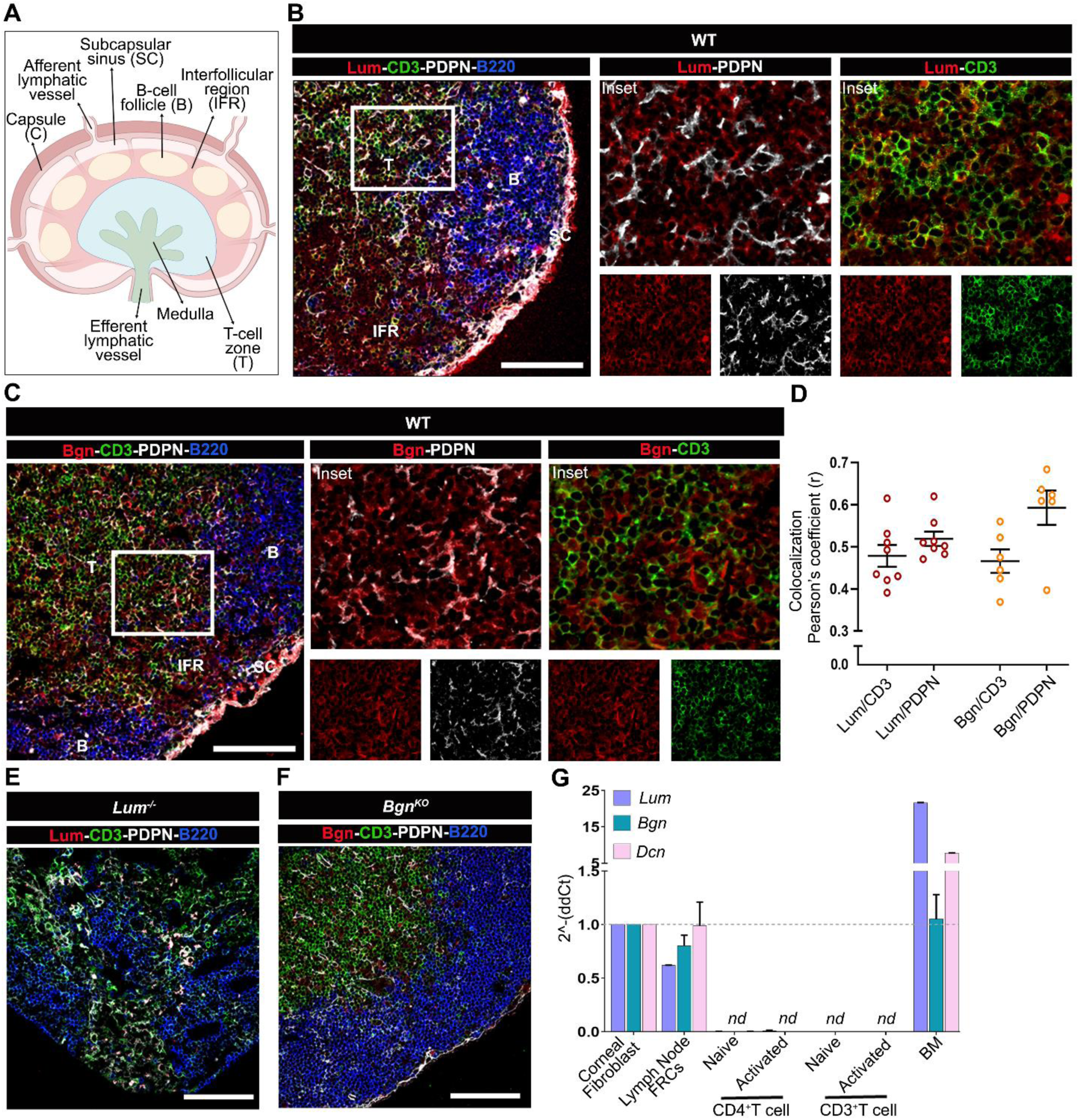
Lumican and biglycan in the lymph node is primarily secreted by stromal FRCs. (**A**) Schematic showing lymph node structure with different sub-regions. (**B** and **C**) Representative confocal images showing localization of lumican or biglycan with podoplanin (PDPN), B220, and CD3 in the lymph node regions of WT mice. Inset shows colocalization of lumican and biglycan with PDPN and CD3. B-B cell follicle, SC-subcapsular region, IFR-interfollicular region, T-T cell zone. (**D**) The cumulative data (n = 6 – 8 images) shows colocalization quantification of images in (B and C) by Pearson’s correlation from 3 independent experiments. (**E** and **F**) Representative confocal images showing the staining specificity of anti-lumican or anti-biglycan antibodies in the lymph node regions of *Lum^-/-^* or *Bgn^KO^* mice. (**G**) *Lum*, *Bgn* and *Dcn* gene expression analyzed by qRT-PCR in WT corneal fibroblast, FRCs from the lymph node of WT mice, naïve and activated CD4^+^ and CD3^+^ T cells from WT mice, and WT bone marrow (BM) cells (n = 2 independent experiments). Scale bars, 100 µm (B, C, E and F). The error bars represent mean ± SEM*, nd = not detected.* See also figures S9 and S10.

Given the colocalization of lumican and biglycan with PDPN and CD3, we next tested expression of *Lum*, *Bgn*, and *Dcn* by FRCs and T cells. Our qRT-PCR data shows that naïve and anti-CD3/anti-CD28 activated CD4^+^ T cells isolated from WT mouse spleens and LNs do not express lumican, biglycan, or decorin. But the CD45^-^CD31^-^PDPN^+^ FRCs from WT LNs expressed all three SLRPs. As these proteoglycans are predominantly expressed in the cornea, we used WT mice corneal fibroblasts as a positive control **(Fig. 7G)**.

## Discussion

Our study shows that contact dermatitis is exacerbated in mice lacking the ECM proteoglycans, lumican, biglycan or decorin. While these SLRPs are known to regulate innate inflammatory signals, the results presented here speak of a novel paracrine regulation of CD4^+^ T cell activation and proliferation that underpin protection against hapten-mediated adaptive immune response. SLRPs are prominent in the ECM of connective tissues. Lumican and biglycan are present in the ear pinnae dermal ECM, and increase after a DNFB challenge. Biglycan also shows a dramatic increase in the epithelial layers of the inflamed ear pinna. We show specific location of all three SLRPs in LNs, in the T cell zone and interfollicular region, and thus ideally situated to regulate T cell functions in CHS. They colocalize with PDPN in the dLN, especially biglycan and decorin, indicating their presence in FRCs in the T cell zone. This is consistent with an earlier report of their expression and location in ER-TR7^+^ FRCs around the conduit core of LNs (*5, 47, 48*). Our qRT-PCR data also shows that *Lum*, *Bgn* and *Dcn* are primarily expressed by the FRCs and not by the naïve or activated CD4^+^ T cells. Indeed, LN FRCs are known to secrete chemokines, cytokines, and lipid mediators with pleiotropic effects on T cell functions (*30*). SLRPs are an important new class of FRC secreted paracrine factor that need to be studied further.

The DNFB-challenged ear pinnae of *Lum^-/-^* and *Bgn^KO^*mice show higher infiltration of CD4^+^ T cells, while their dLN show increased proliferating PCNA^+^CD4^+^ T cells and IFN-γ secreting CD4^+^ T (Th1) cells. Further, activation of the IFN-γ-STAT-1-T-bet signaling axis was evident in *Lum^-/-^* and *Bgn^KO^* mice as dLN lysates from DNFB-mice show elevated p-STAT1. Additionally, in an adoptive transfer model, compared to WTs, the *Lum^-/-^* and *Bgn^KO^* mice show increased activation of OTII cells (CD69^+^CD4^+^) in the dLN after an OVA challenge. Corroboratively, *in vitro* recombinant lumican, biglycan and decorin suppressed T cell activation. ICAM-1 is an important adhesion receptor on antigen presenting cells that interacts with LFA-1 on T cells, to facilitate T cell activation (*49–52*). Interestingly, all three SLRPs interfere with ICAM-1 mediated T cell activation, assessed here as phosphorylation of ZAP-70, Lck and Akt. Their suppressive effects on T cell activation are LFA-1 dependent, as blocking its β2 integrin subunit abrogated lumican, biglycan and decorin functions. Altogether, all three ECM proteoglycans can act as a paracrine signal to restrict CD4^+^ T cell activation by interacting with LFA-1 and thereby inhibit the key signaling pathways downstream of the TCR.

Overall, CHS and ear pinnae pathologies were worse in the SLRP-null strains. The protective functions of SLRPs in CHS may arise from their dual role in early innate immune signals and in adaptive immunity through regulations of T cell functions. In a prior study of 2-4-5, trinitrobenzene sulfonic acid (TNBS) mediated colitis model, the early pro-inflammatory responses from neutrophils and macrophages were low in *Lum^-/-^* mice due to decreased chemotaxis and TLR4 signaling but the overall tissue damage and severity was worse (*53*). In the CHS model however, the *Lum^-/-^*, *Bgn^KO^* and WT mice show comparable infiltration of neutrophils and macrophages in the ear pinnae. This indicates that the elicitation phase is mainly dominated by the adaptive immune cells compared to the early sensitization phase that is driven by the innate immune cells (*4*).

An earlier experimental autoimmune encephalomyelitis (EAE) study reported worse disease in *Lum^-/-^* mice possibly due to increased infiltration of CD4^+^ T cells into the central nervous system (*54*). While we observed a similar increase in pathogenic CD4^+^ T cells into the null mouse skin tissue during CHS, our study differs in two key ways. First, the infiltrating CD4^+^ T cells in CHS are primarily IFN-γ producing Th1 subsets whereas the EAE study reported elevated IL-17 producing Th17 cells. Albeit, the two models have inherent differences in CD4^+^ T cell biases: the CHS model favors a Th1, while EAE harbors Th17 cells. Second, in our study, lumican and biglycan interact with LFA-1 to restrain CD4^+^ T cell proliferation and activation, with little to no effect on CD8^+^ T cells. In the EAE model, the increase in Th17 cells in the *Lum^-/-^*mice was attributed to increased proliferation and decreased apoptosis. Interestingly, many studies have considered close regulation of cell proliferation and apoptosis by these SLRPs, although this has not been tested in T cells. For example, decorin is known to suppress macrophage and tumor cell proliferation through the induction of cyclin-dependent kinase inhibitors, p21 and p27 (*55*). While lumican suppresses proliferation of fibroblasts through p21, and apoptosis through Fas and FasL (*56*).

Biglycan behaves as an endogenous ligand for TLR2 and TLR4, and promotes OVA-specific activation of MHC-I-restricted OT-I and MHC-II-restricted OT-II cells, indicating an increase in DC-T cell interaction. While in a myocardial infarction model, the *Bgn^KO^* mice infiltration of CD3^+^ T cells into the myocardium was decreased, where the underlying cause was attributed to sub-optimal DC-T cell interactions (*57*). By contrast, our CHS study reveals a direct role for lumican and biglycan in LFA-1 – mediated regulation of T cell proliferation and activation. This paracrine regulation is crucial to prevent excessive immune responses and tissue damage that could occur with uncontrolled Th1 activation. Overall, this indicates that binding of SLRPs to LFA-1 possibly disrupts the actin-cytoskeletal dynamics, destabilizing the DC-T cell interaction and formation of proper IS. Our data show a low level of phosphorylated ZAP-70, which confirms that lumican and biglycan destabilize the IS by binding to LFA-1 and decrease the downstream TCR signaling and T cell activation. LFA-1 binding with ICAM-1 also induces specific intracellular signals called “outside-in signals” that lower the threshold dose of antigen required to activate the T cell fully (*45, 58*). Although it remains to be tested, this may be another interesting mechanism by which SLRPs can restrain naïve T cell activation by increasing the antigenic threshold.

### Limitations of the study

While our study investigated the direct involvement of lumican, biglycan and decorin in regulating CD4^+^ T cell activation and proliferation, we have yet to examine the molecular interactions between these SLRPs and LFA-1 and ICAM-1 on the T cell surface. Our data shows a significant increase in the CD8^+^ T cell pool in the dLN of *Lum^-/-^*and *Bgn^KO^* mice but no difference in proliferation compared to WT. We have not tested the activation and effector functions of CD8^+^ cells in the null mice. It will be interesting to investigate these functions in our future studies. Lastly, this study is also limited to SLRP and LFA-1 axis of CD4^+^ T cell regulation and overlooks other possible interactions between SLRPs and T cell surface receptors.

## Materials and Methods

### Mice

All protocols were approved by the New York University Institutional Animal Care and Use Committee (IACUC). All mouse strains were in the C57BL/6J background. Breeding pairs of C57BL/6J (Stock#000664) and *Dcn^-/-^* (Stock#027672) were obtained from the Jackson Laboratories. OT-II (*36*), *Bgn^-/-^* and *Bgn^-/0^* (*13*) and *Lum^-/-^* (*12*) and *Dcn^-/-^*were used to maintain the colonies in a specific pathogen-free mouse facility at New York University, School of Medicine. The mice were housed in clear, air-filtered cages with 12-hour light/dark cycle and ad lib feeding. Mice, between the ages of 8-16 weeks, were used in all experiments, and the animals were sex-balanced within each experiment. The contact hypersensitivity model and OT-II adoptive transfer model were approved by the New York University Institutional Animal Care and Use Committee (IACUC). All CHS ear measurements was blinded. No animals were excluded unless they were sick or with skin injury prior to CHS and OT-II transfer model.

### Contact hypersensitivity (CHS) model

We adapted the CHS model to assess the cell-mediated immune function in the skin during delayed-type hypersensitivity (DTH) due to exogenous hapten exposure (*2*). Briefly, wild-type (WT), *Lum^-/-^* and *Bgn^-/-^* and *Bgn^-/0^* mice were sensitized with 20 μl of 0.5% 2, 4-dinitrofluorobenzene (DNFB; Sigma-Aldrich, St. Louis, MO), in acetone and olive oil (4:1), on shaved abdominal skin day 0 and day 1. At day 5, CHS was elicited by applying 20 μl of 0.2% DNFB on the ear. The unchallenged mice ear was treated with acetone/olive oil alone. Ear thickness was measured in a blinded manner with a digital Vernier caliper (Peacock, Japan) after 24 and 48 hours of the challenge. The degree of CHS was determined as swelling of the DNFB-challenged ear compared with that of the vehicle-treated ear and was expressed in mm. On day 2 post-challenge, animals were euthanized to harvest the ear tissue and the auricular draining lymph node was harvested for further analysis and histological studies.

### OT-II adoptive transfer model

Naïve CD4^+^ T cells from OT-II mice were isolated as previously described with modifications (*37*). Briefly, CD4^+^ T were isolated from the spleen and LN of OT-II TCR Tg GFP^+^ mice using the Easysep Mouse CD4^+^ T cell isolation kit (Stemcell Technologies) or MojoSort Mouse CD4 Naïve T cell isolation kit (BioLegend) according to the manufacturer’s instruction. Generally, on day 0, 1×10^6^ OT-II cells were transferred into the recipient (WT, *Lum^-/-^*, *Bgn^KO^, Dcn^-/-^*) mice by retro-orbital injection in 200 μl. On day 1, 10 μl emulsion of 1:1 IFA and 25 μg OVA_323-339_ peptide (OVA) (Sigma) in PBS was injected into the right hind footpads (s.c.) after 24 hours of T cell transfer. On day 2, animals were euthanized and the popliteal dLN were harvested for further analysis. Single-cell suspension from popliteal LN was stained for live-dead cells, CD4^+^ T cells, and the activation marker, CD69, and were analyzed by flow cytometry.

### Isolation of cells from secondary lymphoid organs and ear tissue

CD4^+^ T cells were isolated from the spleen and LN of WT mice by mechanical disruption, filtered through a 70 μm cell strainer, and by negative selection through the magnetic column using the Easysep Mouse CD4^+^ T cell isolation kit or MojoSort Mouse CD4 Naïve T cell isolation kit. Cells from the ear tissue were isolated as previously described (*37*). Briefly, ears were peeled into two halves and minced using scissors and forceps. Tissues were then incubated for 1.5 hours at 37°C in an orbital shaker (200 rpm) with 1 ml of digestion media containing 0.25 mg/ml Liberase TL (Roche) and 0.25 mg/ml DNase I (Sigma) in RPMI 1640 supplemented with 50 mM β-mercaptoethanol (BME) and 20 mM HEPES. The digestion mix was then inactivated with 1 ml of ice-cold RPMI 1640 containing 10% FBS and 1 mM EDTA. Digested ear tissue was further homogenized using the tissue homogenizer (Qiagen, TissueLyser LT) for 1 minute. The cells were then passed through a 70 μm cell strainer. Live cells were counted by trypan blue exclusion dye (Sigma) and cells were further used for flow-cytometry staining.

### In vitro restimulation of T cells

For flow cytometric analysis of intracellular cytokines, IFN-γ, IL-17 and IL-4 in CD4^+^ T cells from dLN of CHS mice were restimulated with PMA and ionomycin and Brefeldin-A. Lymphocytes from ear dLN of DNFB-challenged and unchallenged (WT, *Lum^-/-^* and *Bgn^-/-^*) mice were isolated by mechanical disruption, filtered through a 70 μm cell strainer, followed by RBC lysis and cells were finally resuspended in RPMI1640 medium containing 50 mM BME, 10 mM HEPES, 1x antibiotic/antimycotic and 1% charcoal-stripped serum (Gibco). Cells were stimulated for 5 hours at 37°C with PMA (50 ng/ml) (Sigma) and ionomycin (420 μg/ml) (Sigma) to induce IFN-γ, IL-17 and IL-4 production. Cells were treated with 1x Brefeldin-A (5 mg/ml) for the last 3 hours. After 5 hours, the cells were washed with ice-cold 1x PBS and stained for intracellular cytokines.

### Isolation and sorting of lymph node fibroblastic reticular cells (FRCs)

Lymph nodes from WT mice were enzymatically digested using a previously published protocol (*59*). Briefly, inguinal, popliteal, cervical, axillary, brachial and mesenteric lymph nodes from 3 mice/genotype were pooled and cut into pieces and digested with the 1 ml of enzyme mix containing 0.8 mg/ml Dispase (Roche), 0.2 mg/ml Collagenase P (Roche) and 0.2 mg/ml DNase I (Sigma) in RPMI 1640. Tissue with the enzyme mix was incubated at 37°C in a water bath and gently rocked at every 5 min intervals to mix the contents. After 20 minutes, LN pieces were gently triturated with a 1 ml pipette to disrupt the capsule. The LN fragments were allowed to settle for the 30s, and the enzyme mix was carefully removed and added to a 15ml tube with complete RPMI 1640. 2 ml of fresh enzyme mix was added to the fragments and incubated back in water bath with gentle trituration at 5 minutes intervals. After every 10 minutes, the enzyme mix was collected and the fragments were mixed with fresh enzyme mix. This process was repeated for a total of 60 minutes. At 1 hour, all the collected supernatant was centrifuged (300 g, 4 min, 4°C) and resuspended in ice-cold FACS buffer (2% FBS, 5 mM EDTA in PBS), filtered through 100 μm cell strainer and enriched for CD45^-^ cells by removing the CD45^+^ cells using the MojoSort mouse CD45 nanobeads kit (Biolegend). Cells were then stained with Live-Dead Blue, CD45, CD31, and PDPN and sorted for the FRCs (CD45^-^CD31^-^PDPN^+^) using the Sony SH800z sorter (Sony) using 100 µm chip and 20 psi stream pressure. Viability was 80%, and the purity of sorted FRCs was 90%. FRCs were directly used for RNA extraction.

### In vitro CD4^+^ T cell activation assays

Naïve CD4^+^ T cells were isolated from WT splenocytes and LN by magnetic negative selection (Easysep Mouse CD4^+^ T cell isolation kit). For the in vitro activation assay, 96 well flat bottom plates were coated with or without anti-CD3/CD28 antibodies for O/N at 4°C. The next day, plates were washed 2x with warm 1x PBS and coated with 2.5 μg/ml of recombinant (r) mouse/human lumican (rLum) or biglycan (rBgn) or decorin (rDcn) for 2 hours at room temperature (RT), followed by washes and naïve CD4^+^ T cells were added and cultured for 6 hours for early activation (CD69) and 24 hours for late activation (CD25) and analyzed by flow cytometry. 2.5 μg/ml of rLum, rBgn, and rDcn were consistently used for all the experiments unless otherwise mentioned.

### CD4^+^ T cell chemotaxis assay

Isolated naïve CD4^+^ T cells were either treated with PBS or with 1 µg/ml of rLum, rBgn, and rDcn for 2 hours followed by 2x washes and added onto the upper chamber containing a polycarbonate transwell membrane filter (5-μm pore size; Corning). The lower chamber contained 0.5 µg/ml of rCCL19 in complete RPMI media supplemented with 1% charcoal-stripped serum. After 4 hours, cells migrated to the lower chamber were recovered, stained with trypan-blue exclusion dye for dead cells, and live cells were counted using a hemocytometer.

### Flow cytometry

Lymphocytes were washed with ice-cold PBS and stained with Live-Dead blue, cells were washed twice followed by surface staining. Cells were stained in a 96-well V-bottom polypropylene plate. Cells were blocked with CD16/CD32 Fc block (1:200) in FACS buffer (PBS with 2% FBS and 5mM EDTA) at 4°C, washed by centrifugation at 1900 rpm, 3 minutes at 4°C. For surface staining, cells were stained with fluorophore conjugated antibodies, washed 3x with FACS buffer, fixed in 4% paraformaldehyde, and resuspended in FACS buffer for analysis.

For intracellular staining, fixed cells were permeabilized by resuspended in 1x Intracellular staining perm wash buffer (BioLegend) and centrifuged at 350 g, 10 minutes at RT. This was repeated 2 times followed by staining with fluorophore-conjugated antibodies against IFN-γ, IL-17 and IL-4 for 60 minutes in the dark at RT. Cells were then washed with the intracellular staining perm wash buffer and resuspended in FACS buffer for analysis. Fluorescence minus one (FMO) is used as a negative control.

For nuclear T-bet staining, after live-dead and surface staining, cells were fixed using True-Nuclear 1x Fix concentrate (BioLegend) in the dark for 60 minutes at 4°C. Cells were washed by centrifugation at 400 g at RT for 5 minutes and resuspended in True-Nuclear 1x Perm buffer. Cells were again centrifuged 2 times at 400 g for 5 minutes and stained with PE-conjugated anti-T-bet antibody diluted in True-Nuclear 1x Perm buffer for 60 minutes. Followed by 3x washes with True-Nuclear 1x Perm buffer and finally resuspended in FACS buffer for analysis.

For Phospho flow staining, cells were washed after treatment and immediately fixed by warm 4% PFA for 15 minutes at RT. Cells were washed 2x with ice-cold FACS buffer and resuspended well in pre-chilled True-Phos Perm buffer (BioLegend) by vortexing. Incubate cells at −20°C for 60 minutes and wash 2x with FACS buffer, followed by staining against phospho-ZAP-70, phospho-Lck and phospho-Akt-1 for 60 minutes, washed 3x, resuspended and analyzed.

Multicolor flow cytometric analysis was performed on flow ZE5 (Yeti) analyzer (Bio-Rad) and analyzed using the FlowJo software (Tree Star, Inc.). All primary conjugated antibodies were used at 1:200 dilution and the antibody details can be found in the **Table S1**.

### In vitro blocking assay

Naïve CD4^+^ T cells isolated from WT mice spleen and LN were treated with or without 10 µg/ml anti-β2 integrin non-function blocking antibody (BioLegend) for 1 hour and added onto 96 well plate coated with anti-CD3/CD28 along with either 2.5µg/ml of rLum or rBgn or rDcn or 1µg/ml of ICAM-1 for 30 minutes to check phospho-Akt-1 expression or 6 hours to check activation CD69 expression or 72 hours to check proliferation (CD4^+^ T cells were pre-labeled with CFSE for proliferation assay). The anti-Rat IgG2a antibody (10 µg/ml) pretreated CD4^+^ T cells served as an isotype control for the anti-β2 integrin blocking antibody. CD4^+^ T cells were harvested at indicated time points, appropriately stained as described earlier, and analyzed on a ZE5 analyzer (BioRad).

### Immunofluorescence staining

Naïve CD4^+^ T cells isolated from WT mice spleen and LN were plated in 8-chamber slides coated with poly-L-Lysine. Cells were then treated with or without anti-CD3/CD28 along with rLum or rBgn tagged with 6X HIS for 60 minutes. Cells were washed with PBS followed by fixation (4% PFA) for 10 min, RT and blocked with 1% BSA in PBS for 1 hour at RT, then incubated with anti-CD18 and AF 488 conjugated anti-HIS antibodies for ON at 4oC in the dark. Cells were washed and incubated with AF 555 conjugated anti-rabbit antibody for 2 hours at RT in the dark. Cells were washed and counter-stained with DAPI (nuclear stain), then washed and sealed in mounting media with 1.5 mm coverslips. Slides were imaged using a Zeiss LSM 880 confocal microscope. The histograms for the fluorescence intensities of the merged panels were analyzed using the Plot profile tool of Fiji (Image J) software.

### Immunohistochemistry and histology

For H&E staining, ear pinnae were excised at 48 hours after the DNFB challenge and fixed in 10% formalin, paraffin embedded, cut into 7 µm thick cross-sections and stained with routine hematoxylin and eosin (H&E). To examine the infiltration of cells, images of H&E stained ear tissue cross sections were captured with a Nikon Eclipse E400 microscope fitted with a DXM1200 Nikon Digital camera.

For multiplex immunofluorescence and imaging of ear tissue, 5 µm formalin fixed paraffin-embedded sections were stained with Akoya Biosciences Opal multiplex automation kit reagents (Leica Cat#ARD1001EA) on a Leice BondRX autostainer, according to manufacturer’s instructions. Briefly, slides were treated with epitope retrieval with Leica Biosystems Epitope Retrieval 2 solution followed by primary and secondary antibody incubation and tyramide signal amplification with Opal fluorophores as indicated in Supplementary Table 1. Primary and secondary antibodies were removed during sequential epitope retrieval, but the relevant fluorophores remained covalently attached to the antigen. Slides were counterstained with spectral DAPI (Akoya Biosciences, FP1490) and mounted with ProLong Gold Antifade (ThermoFisher). Semi-automated image acquisition was performed on a Vectra Polaris multispectral imaging system. Whole slide was scanned at 20x in the motif mode and the region of interest was selected for spectral unmixing and image processing using InForm v.2.4.10 software from Akoya Biosciences. Infiltrating cells were counted per one field of view. Five fields of view were selected at random in one section, and slides from two mice were examined for each staining. Unchallenged and challenged ear tissue images were unlabeled and mixed, and cells were counted manually and blindfolded in an unbiased manner by two researchers.

For OCT frozen tissues, cryosections were air-dried for 2 h, fixed in 4% PFA for 10 minutes, and blocked for 1 h with 5% normal donkey or goat serum in 1x PBS with 3% BSA. Sections were then incubated O/N at 4°C with primary antibodies diluted in 1x PBS with 0.05% Tween-20 (PBST) at a predetermined optimal concentration. No primary antibody-treated sections were used as a control for nonspecific staining. This is followed by washing with PBST and staining with secondary antibodies diluted in a blocking buffer for 1 h at RT. Followed by 3 times washing with PBST and counter-stained with 1 µg/ml DAPI for 2 min. Slides were washed twice with PBS and mounted in antifade mounting media with a coverslip. Slides were imaged using Zeiss LSM 880 confocal microscope and images analyzed using Fiji. To quantify colocalization, analysis was done on three separate planes for each field using Fiji and the Coloc2 plug-in was used to generate a binary image of colocalized pixels from two separate channels and background pixel intensity was eliminated using Costes automatic threshold method from all images and Coloc2 was run for the field. The colocalization was plotted as Pearson’s coefficient (r).

### Immunoblotting and densitometry analysis

Lymphocytes from the ear dLN of unchallenged and DNFB-challenged mice were harvested and lysed in lysis buffer (25 mM MES, pH-6.5, 150 mM NaCl, 1% Triton X-100, 1 mM PMSF, 1 mM NaF, 0.1 mM Na3VO4 and 1x PIC) and homogenized using the pre-chilled tissue homogenizer (Qiagen, TissueLyser LT) for 1 minute. The tissue homogenate was centrifuged at 10,000 rpm for 20 minutes at 4°C, and supernatant was collected. Protein concentration was quantified by Coomassie Blue protein assay, and 60 µg of lysates were separated on NuPAGE 4-12%, Bis-Tris gel (Invitrogen) with NuPAGE MOPS SDS running buffer (Invitrogen). The gels were blotted onto PVDF membranes using a semi-dry apparatus, blocked, and incubated with primary antibodies (1:1000): anti-phospho-STAT-1, -STAT-1, and –β-actin (Cell Signaling). Appropriate HRP-linked secondary antibodies (1:5000) were used to detect the primary antibodies. Original uncropped immunoblots can be found in the supplementary data figure S5. Band intensities of phospho-STAT-1 were quantified using Fiji and normalized to total STAT-1 band intensities.

### Quantitative qRT-PCR

Total RNA was extracted from the FACS sorted WT FRCs (CD45^-^ CD31^-^PDPN^+^) cells using RNeasy Mini kit (Qiagen) and cDNA was synthesized using cDNA reverse transcription kit (BioRad). The cDNA (10-20 ng) was used for qRT-PCR using Applied Biosystems TaqMan assays for genes on a One Step Plus instruments (Applied Biosystems). The Ct was normalized against the housekeeping gene β-actin.

### Statistical Analysis

All the experiments were repeated two to four times, as indicated in the figure legends. Data are presented as mean ± SEM, also indicated in each figure legend. Comparisons between multiple groups were performed by one-way or 2-way ANOVA followed by multiple comparisons analysis. Comparisons between the two groups were performed using a Unpaired t test with Welch correction. P ≤ 0.05 is considered to be significant. All statistical analyses were performed using Graph Pad Prism version 10.

## Acknowledgments

We thank Dr. Sergei B. Koralov for his valuable comments. We thank the NYU Langone Microscopy Laboratory (RRID: SCR_017934), Experimental Pathology Research Laboratory (RRID: SCR_017928) and Cytometry and Cell Sorting Laboratory which are partially supported by the Laura and Isaac Perlmutter Cancer Center support grant P30CA016087. We thank Michael Cammer for advice on confocal microscopy and Tansol Choi for helping with preliminary experiments.

## Author contributions

Conceptualization: GM, JF, SC; Methodology: GM, JF, CL; Investigation: GM, JF; Validation: GM, SC; Data Analysis: GM, JF, SC; Supervision: SC, GM; Writing—original draft: GM, SC; Writing—review & editing: GM, JF, CL, SC; Funding: SC

## Data and materials availability

“All data needed to evaluate the conclusion in the paper are available in the main text and the supplementary materials.”

## Supplementary Materials for

**Fig. S1:**
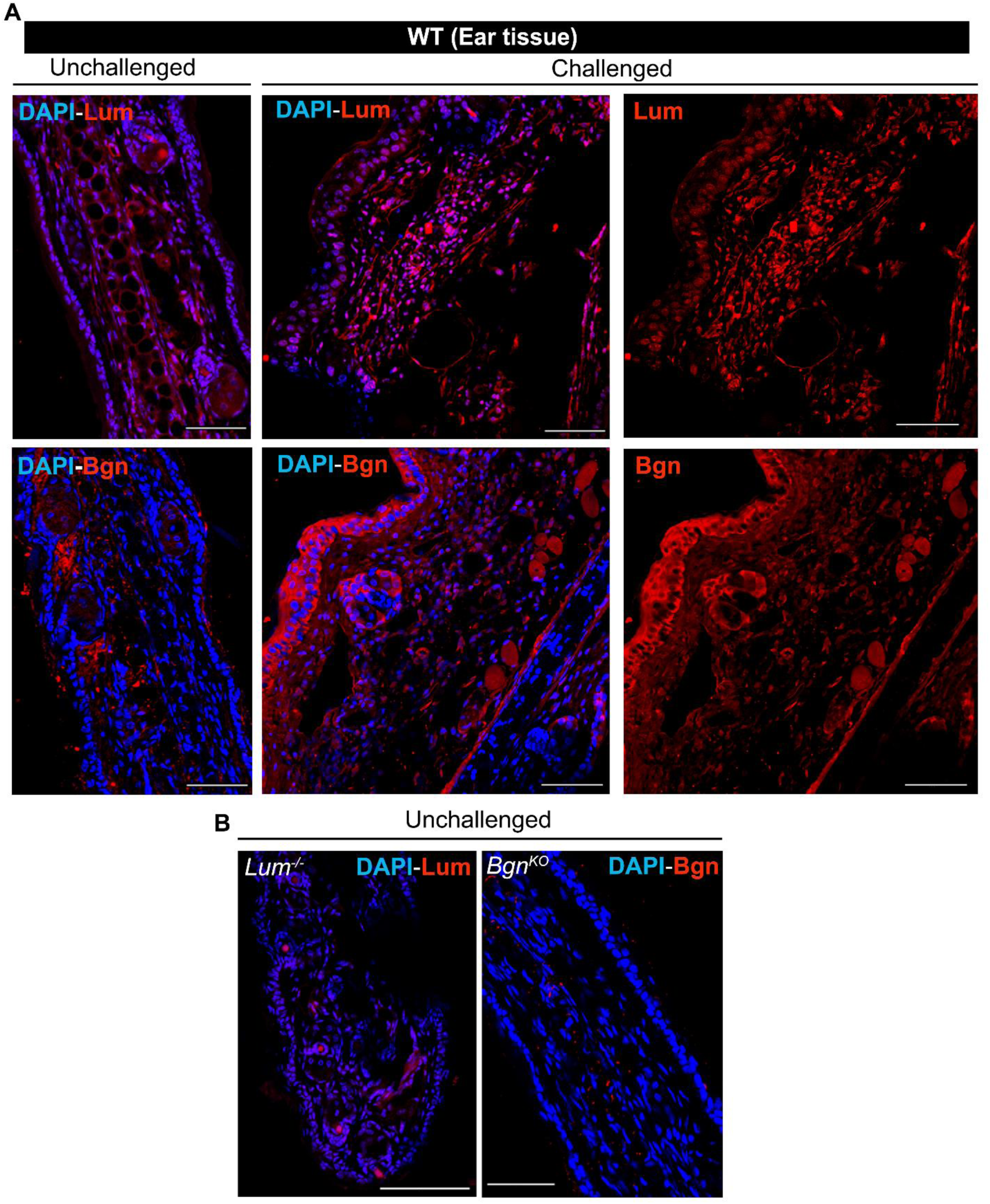
Lumican and biglycan expression in the WT ear tissue. **(A)** Lumican (Lum) and biglycan (Bgn) level increases in the challenged WT ear tissue as shown by the IHC images. DAPI staining indicates the nucleus. **(B)** Ear tissue sections of *Lum^-/-^* and *Bgn^KO^* mouse served as a negative control for the primary antibodies and showed little to no immunoreactivity. Scale bars, 100 µm (A and B).

**Fig. S2:**
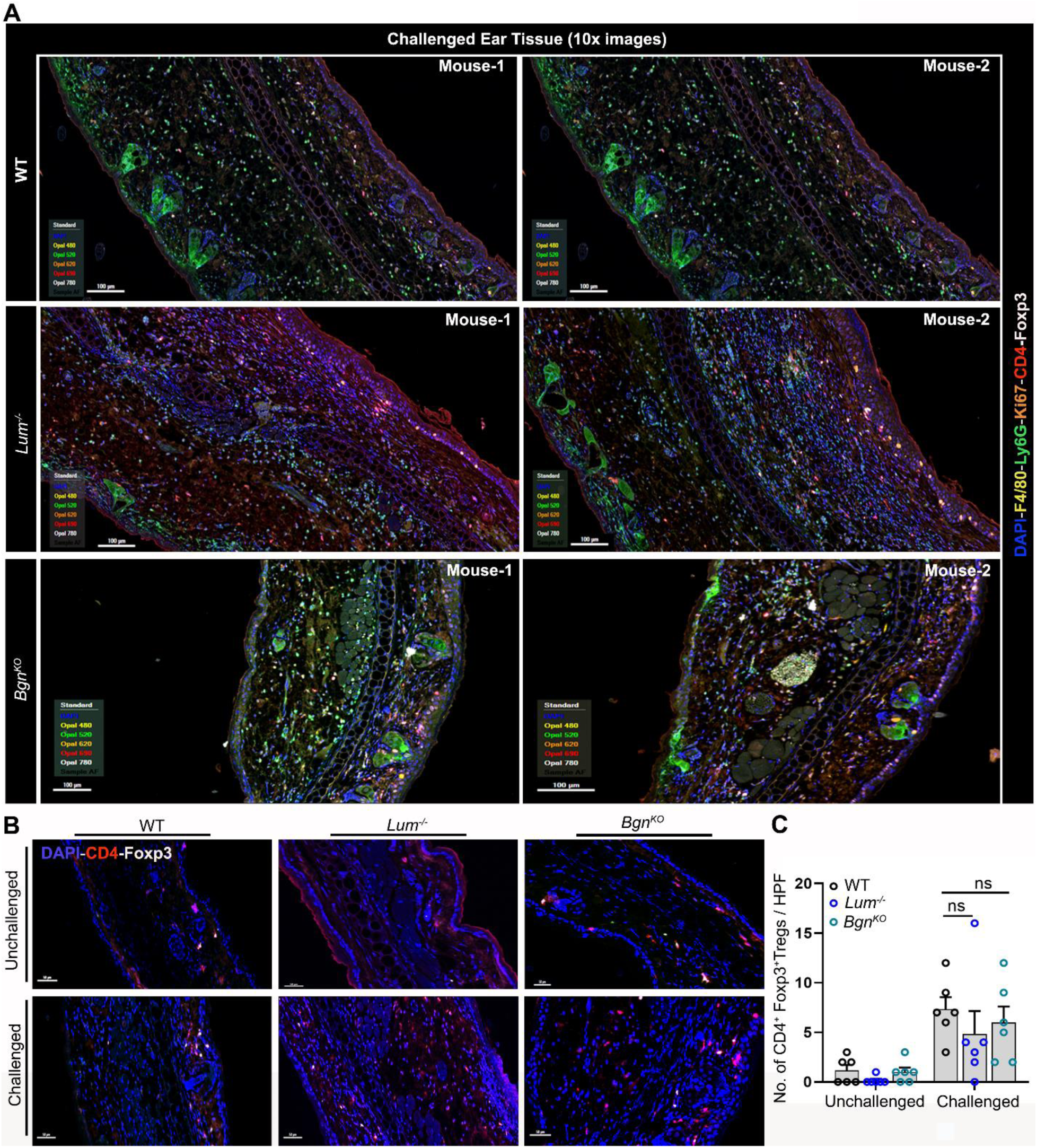
WT, *Lum^-/-^*, *Bgn^KO^* have comparable number of Foxp3^+^ Tregs in the ear tissue. **(A-B)** Wild type and the null mice have similar number of CD4^+^Foxp3^+^ Tregs in the ear tissue as shown by the IHC images (A) and the cumulative bar graph from six high-power field (HPF) from one experiment with two mice per genotype (B). **(C)** IHC images showing all the different markers in the challenged ear tissue at 10x magnification from two mice per genotype. Scale bars, 100 µm (A and C). The error bars represent mean ± SEM, unpaired t test with Welch correction (B). *ns, not significant*.

**Fig. S3:**
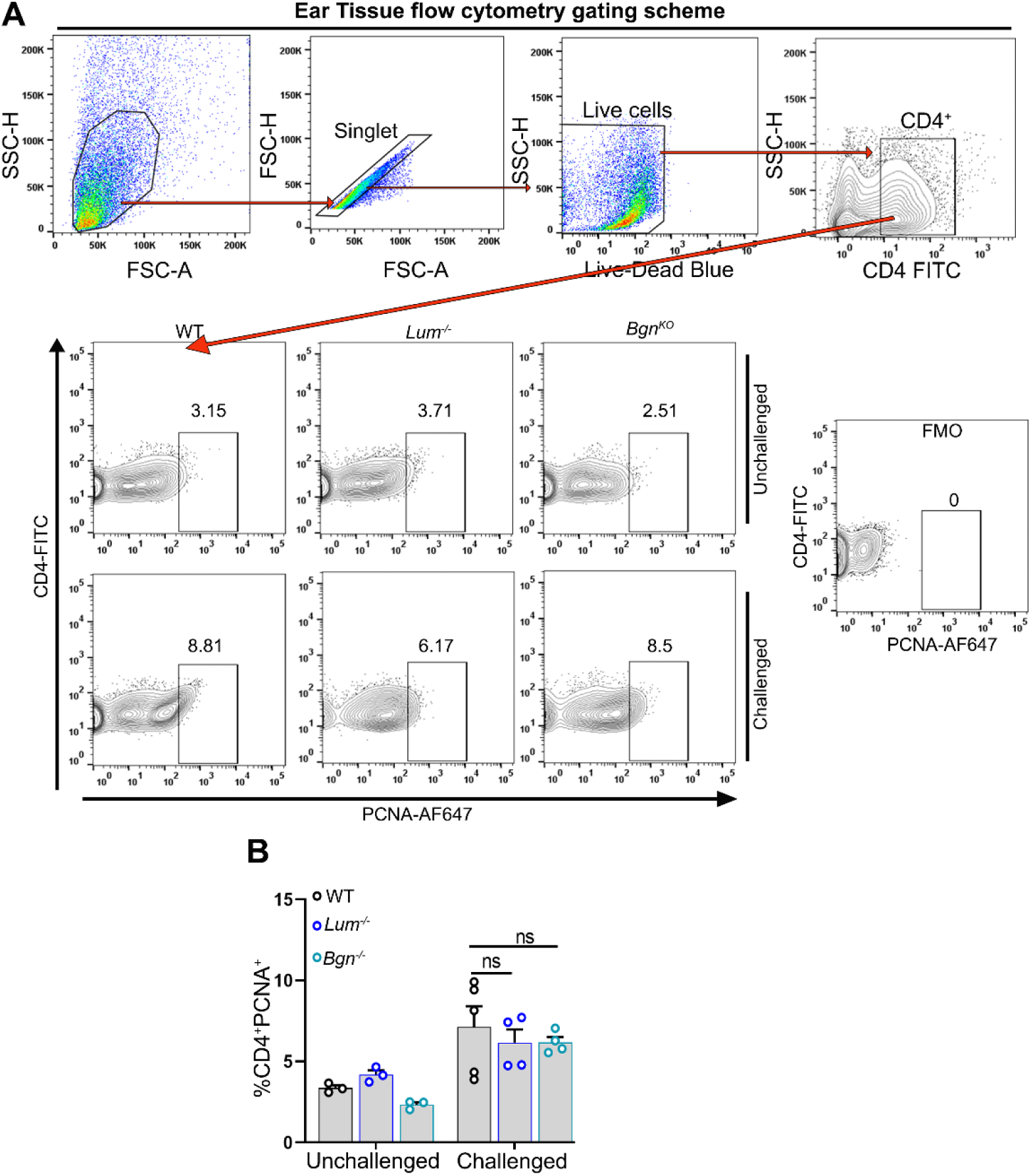
WT, *Lum^-/-^*, *Bgn^KO^* have comparable percentage of proliferating CD4^+^T cells in the ear tissue. **(A)** Flow cytometry gating scheme for the proliferating PCNA^+^CD4^+^ T cells in the ear pinnae. **(B)** Cumulative data from 2 independent experiments (n = 3 – 5 mice/genotypes) shows the percentage of proliferating CD4^+^ T cells (PCNA^+^CD4^+^) in the ear pinnae. The error bars represent mean ± SEM, unpaired t test with Welch correction. *ns, not significant*.

**Fig. S4:**
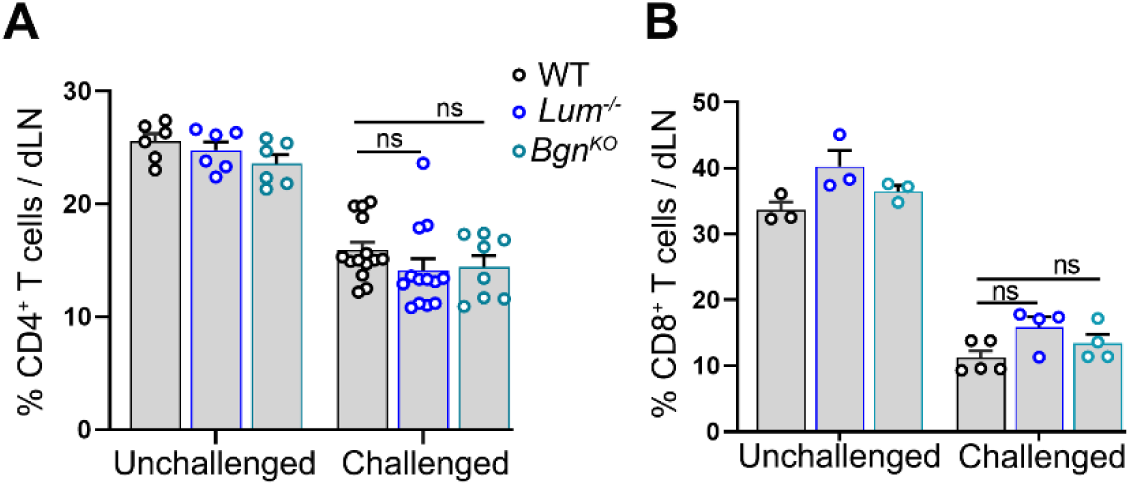
Percentage of CD4^+^ and CD8^+^ T cells in the dLN. **(A)** The cumulative data shows the percentage of CD4^+^ T cells (n = 6 – 14 mice from 3 independent experiments. **(B)** The cumulative data shows percentage of CD8^+^ T cells in the dLN of unchallenged and challenged WT, *Lum^-/-^* and *Bgn^KO^* (n = 3 - 5 mice from 2 independent). The error bars represent mean ± SEM, unpaired t test with Welch correction. *ns, not significant*.

**Fig. S5:**
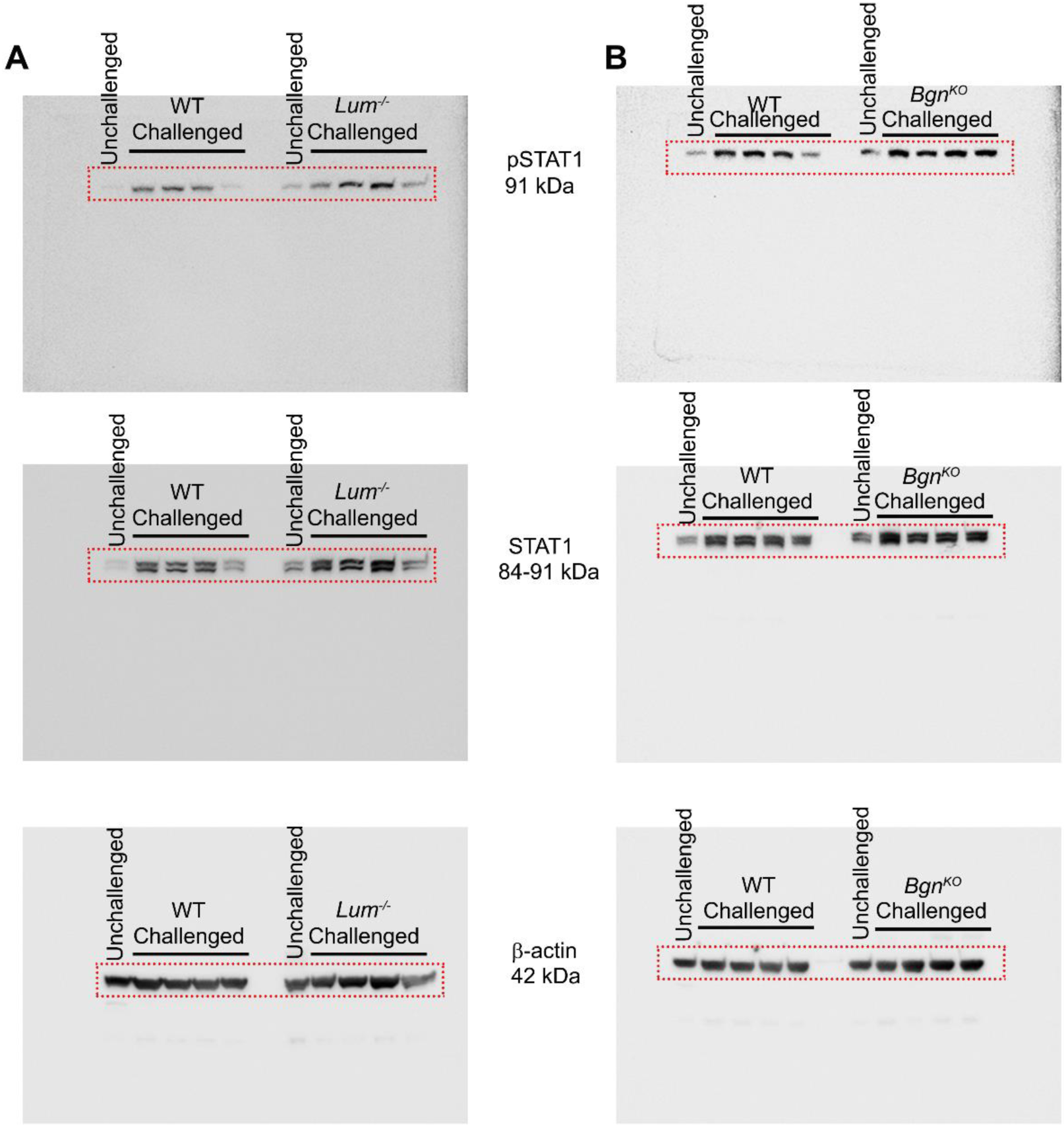
Full length Immunoblots showing phospho-STAT1, STAT1 and β-actin level in the dLN. **(A and B)** Uncropped images of the blots probed for indicated proteins as presented in (A) for main Fig.5D and (B) for main Fig.5E.

**Fig. S6:**
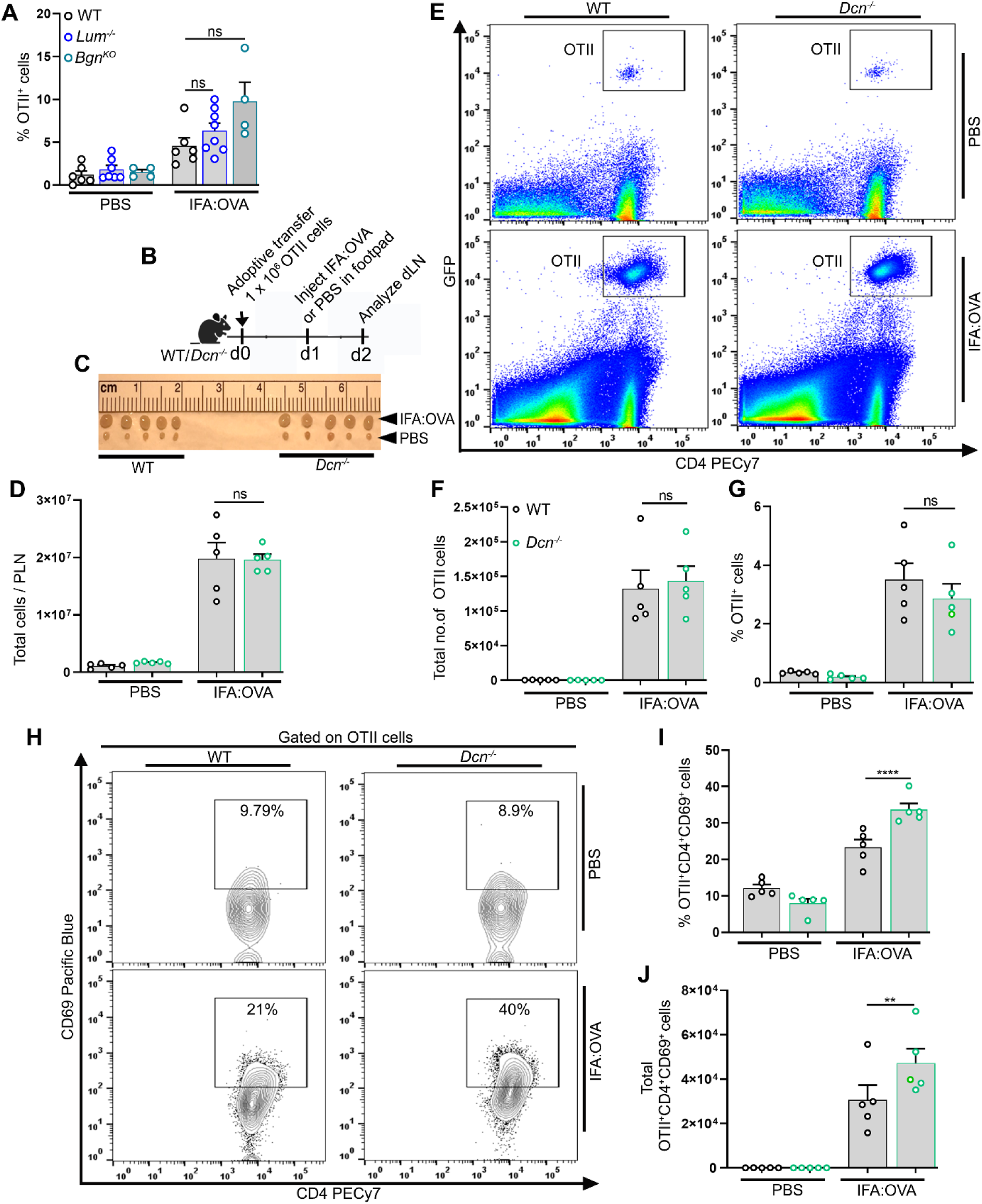
Decorin suppresses CD4^+^ T cell activation *in vivo*. **(A)** Cumulative data show percentage of transferred OT-II cells in the dLN (n = 4 – 8 mice/genotype) from two independent experiments. **(B)** Schematic showing the OT-II adoptive transfer model. On day0, GFP expressing OT-II cells (1 x 10^6^) were adoptive transferred retro-orbitally to WT and *Dcn^-/-^* mice. Foot-pad were challenged with IFA-OVA or PBS on day1 and dLN were harvested for analysis on day2 after euthanasia. **(C and D)** Images showing similar sizes and comparable number of total cells in dLN of WT and *Dcn^-/-^* mice. **(E)** Representative dot plots showing transferred OT-II cells (GFP^+^CD4^+^ cells) in the dLN after injected with either IFA-OVA or PBS in WT and *Dcn^-/-^* mice. **(F and G)** Cumulative data show total number (E) and percentage (F) of transferred OT-II cells in the dLN (n = 5 mice/genotype) from one independent experiment. **(H)** Representative contour plots showing percentage of activated OT-II cells (CD69^+^CD4^+^ cells) in the dLN. **(I and J)** Cumulative data show the percentage (H) and total number (I) of activated OT-II cells (CD69^+^OT-II^+^CD4^+^ cells) in the dLN after injected with either IFA-OVA or PBS in WT and *Dcn^-/-^* mice (n = 5 mice/genotype) from one experiment. The error bars represent mean ± SEM, unpaired t test with Welch correction. \**\*P < 0.01,* \**\*\*\*P < 0.0001*; *ns, not significant*.

**Fig. S7:**
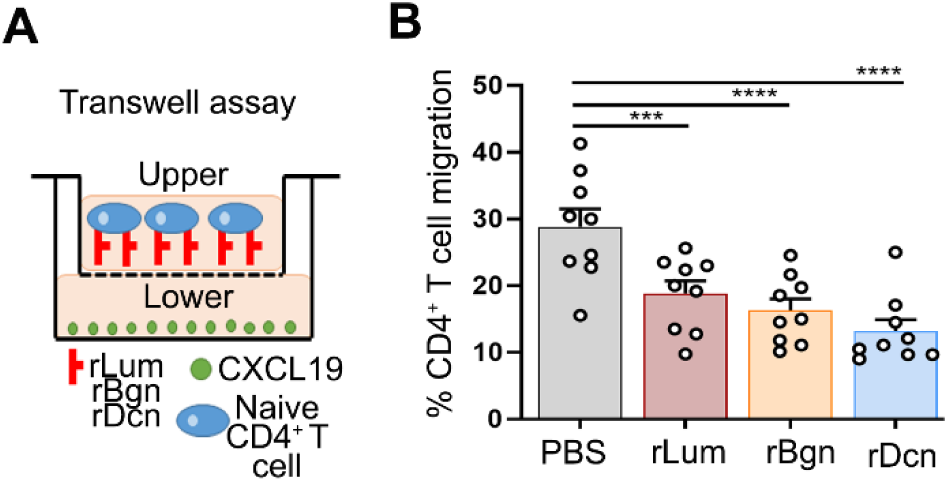
Lumican, biglycan and decorin suppresses CD4^+^ T cell chemotaxis *in vitro*. **(A)** Schematic shows the transwell migration assay. **(B)** Cumulative data from 2 independent experiments shows the number of CD4^+^ T cells treated with 1 µg/ml of either rLum or rBgn or rDcn migrated towards 0.5 µg/ml rCCL19 in the lower chamber as percentage with respect to just media (no rCCL19). The error bars represent mean ± SEM, two-way ANOVA. *\*\*\*P < 0.001*, *\*\*\*\*P < 0.0001*.

**Fig. S8:**
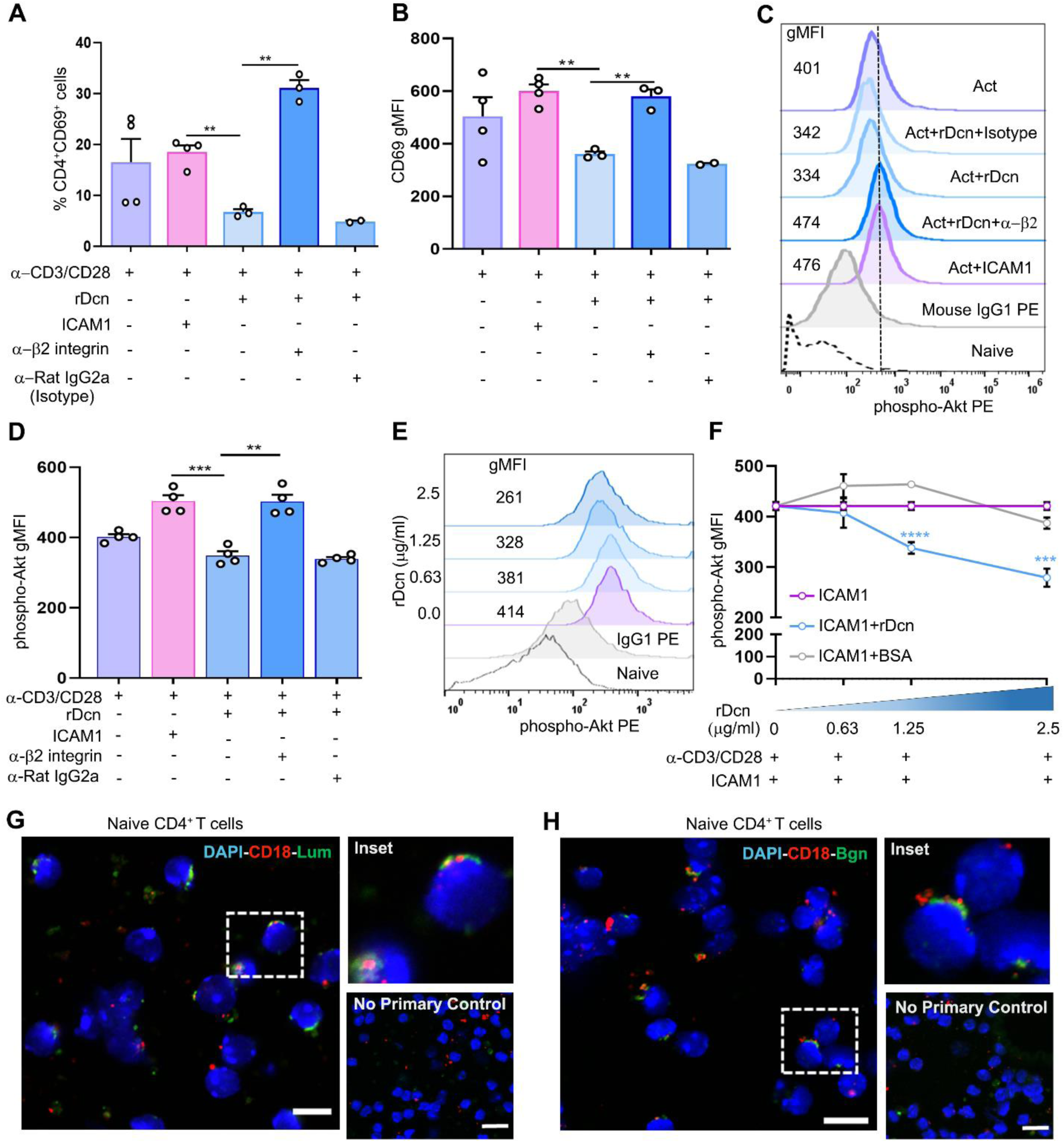
Decorin suppresses CD4^+^ T cell activation. **(A and B)** Cumulative data from 2 independent experiments shows the percentage (A) of activated CD4^+^ T (CD69^+^CD4^+^) cells and surface expression of CD69 (gMFI) (B). **(C)** Representative histograms show the phospho-Akt-1 level in activated CD4^+^ T cells with various treatments as indicated in the plot. **(D)** Cumulative data from 2 independent experiments shows the phospho-Akt-1 levels (gMFI) in activated CD4^+^ T cells with either rDcn or ICAM-1 alone or in combination with anti-β2 integrin antibody. Anti-Rat IgG2a is used as an isotype control. **(E)** Representative histograms show the phospho-Akt-1 level in activated CD4^+^ T cells as an indicative of ICAM-1 binding strength to LFA-1 in the presence of increasing soluble rDcn or BSA. **(F)** Cumulative line graph from 2 independent experiments shows the phospho-Akt-1 levels (gMFI) in activated CD4^+^ T cells as a measure of competitive binding between LFA-1 and rDcn or ICAM-1. (**G** and **H**) Representative confocal image from 2 independent experiments shows staining of CD18 with rLum (G) and rBgn (H) in naïve CD4^+^ T cells. No primary control indicates the specificity of primary antibodies. The histogram depicts cross-line scans of the fluorescence intensities of the merged panels. Scale bars, 10 µm. Each dot represents technical replicates (A, B and D). The error bars represent mean ± SEM, unpaired t test with Welch correction. \**\*P < 0.01,* \**\*\*P < 0.0001,* \**\*\*\*P < 0.0001*; *ns, not significant*.

**Fig. S9:**
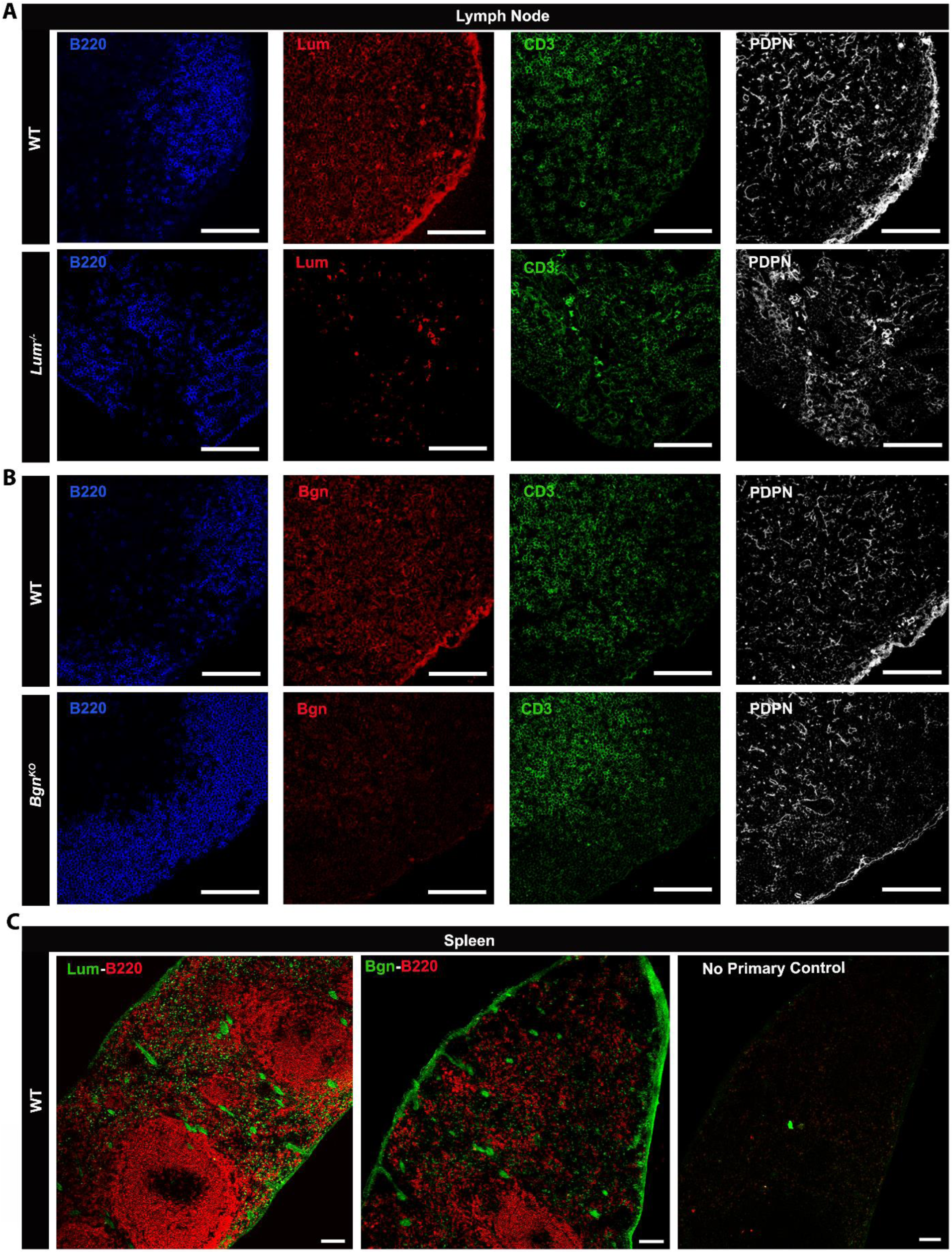
Localization of lumican and biglycan in the LN and spleen. **(A and B)** Confocal images of showing individual channels for the merge images in main Figure 1B and 1C. **(C)** Representative confocal images showing localization of lumican and biglycan, and B220 in the spleen of WT mice. Scale bars, 100 µm (A, B and C).

**Fig. S10:**
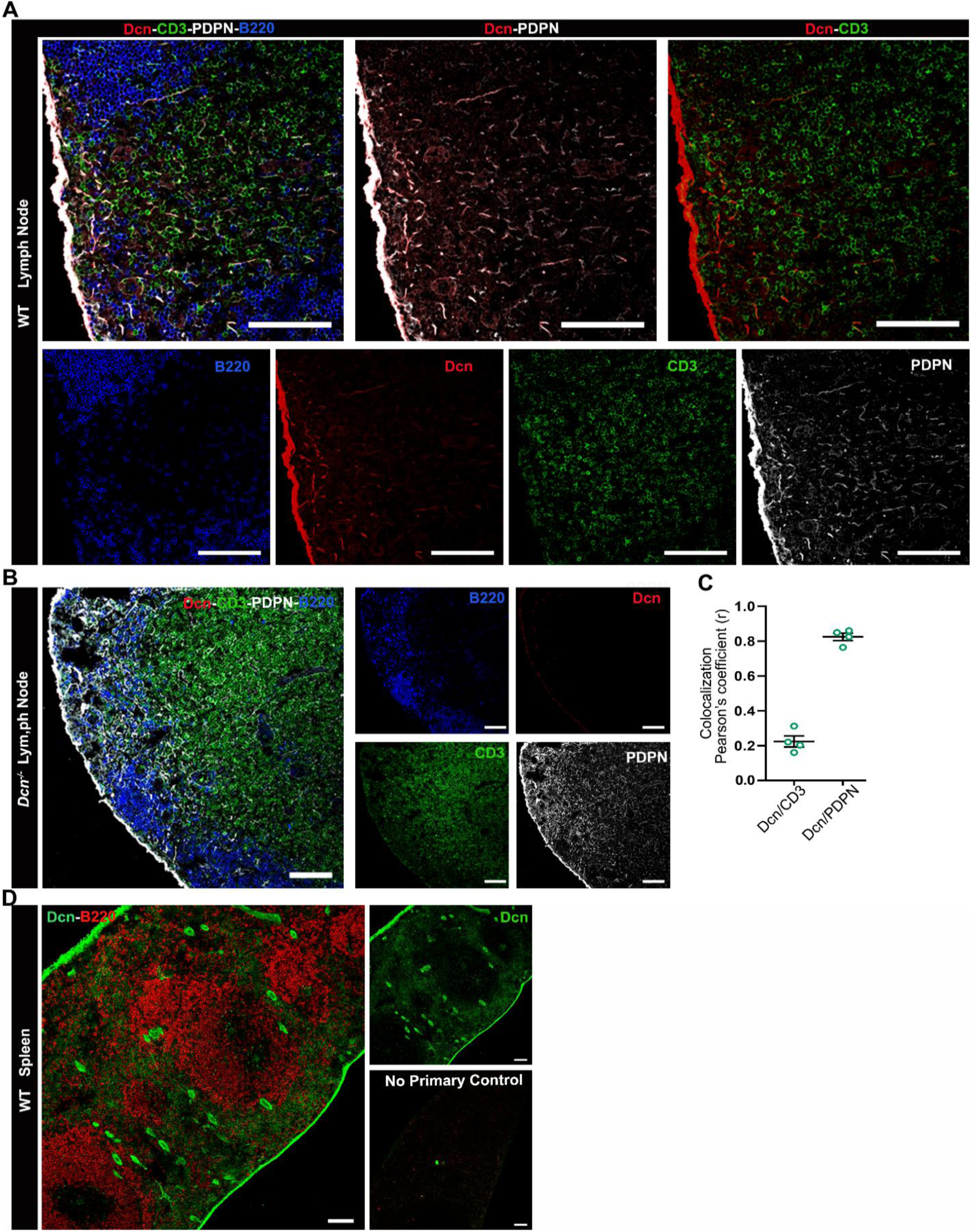
Localization of decorin in the LN and spleen. **(A)** Representative confocal images showing localization of decorin with PDPN, B220 and CD3 in the lymph node regions of WT mice. **(B)** Representative confocal images showing the staining specificity of anti-mouse decorin antibody in the lymph node regions of *Dcn^-/-^* mice. **(C)** The cumulative data (n = 4 images) showing colocalization quantification of images in (A) by Pearson’s correlation from 2 independent experiments. **(D)** Representative confocal images showing localization of decorin and B220 in the spleen of WT mice. Scale bars, 100 µm. The error bars represent mean ± SEM.

**Table S1:**
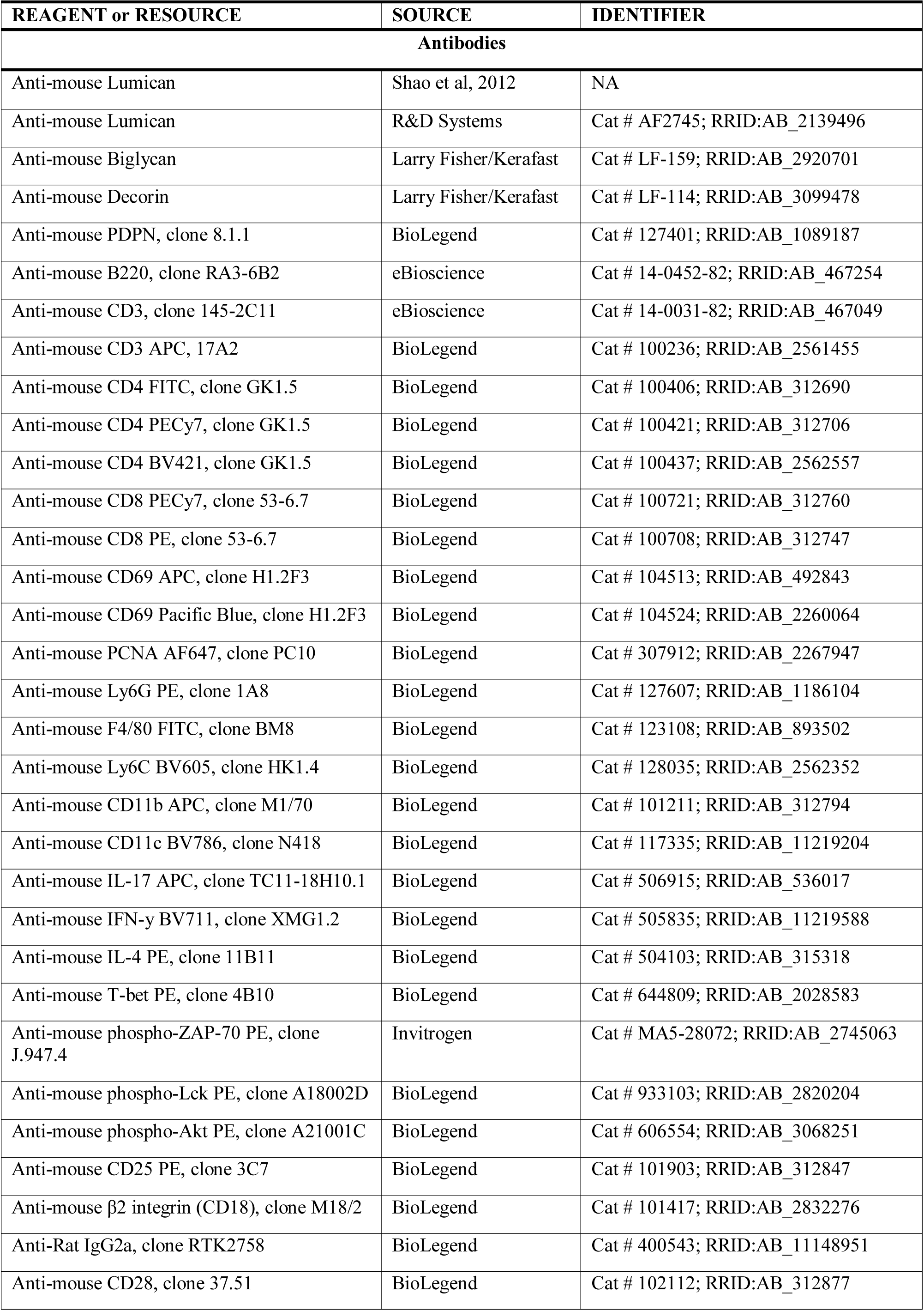
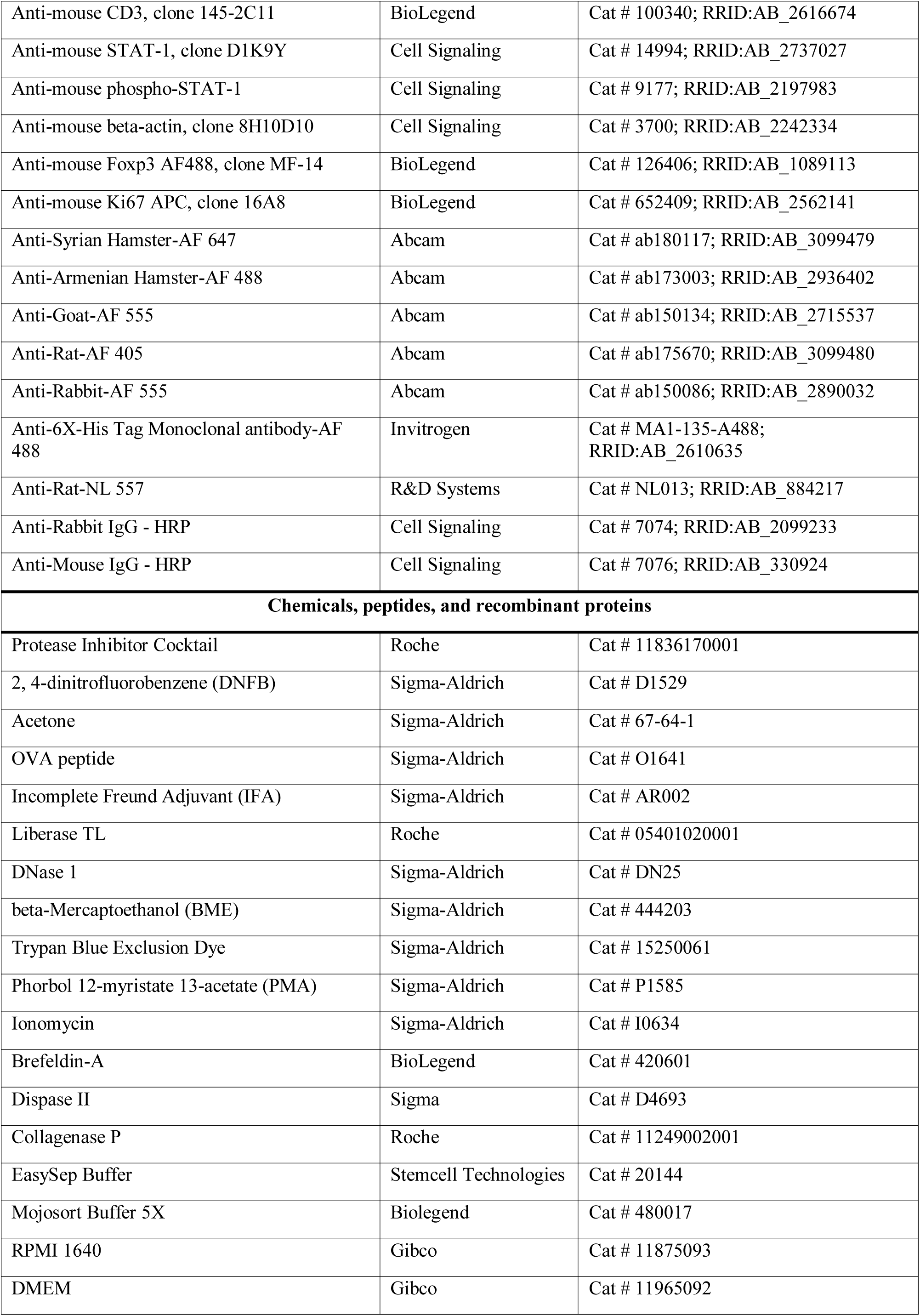
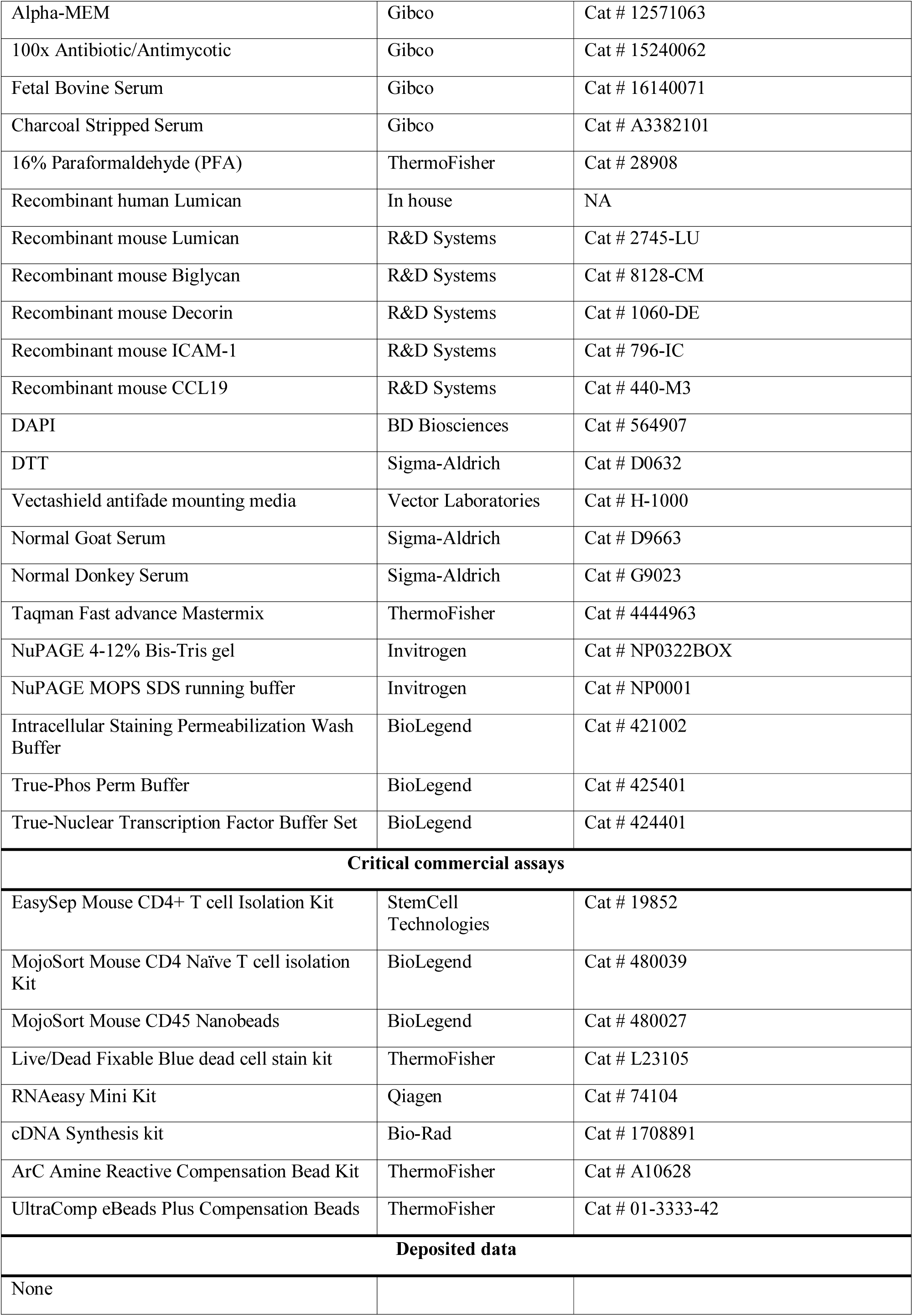
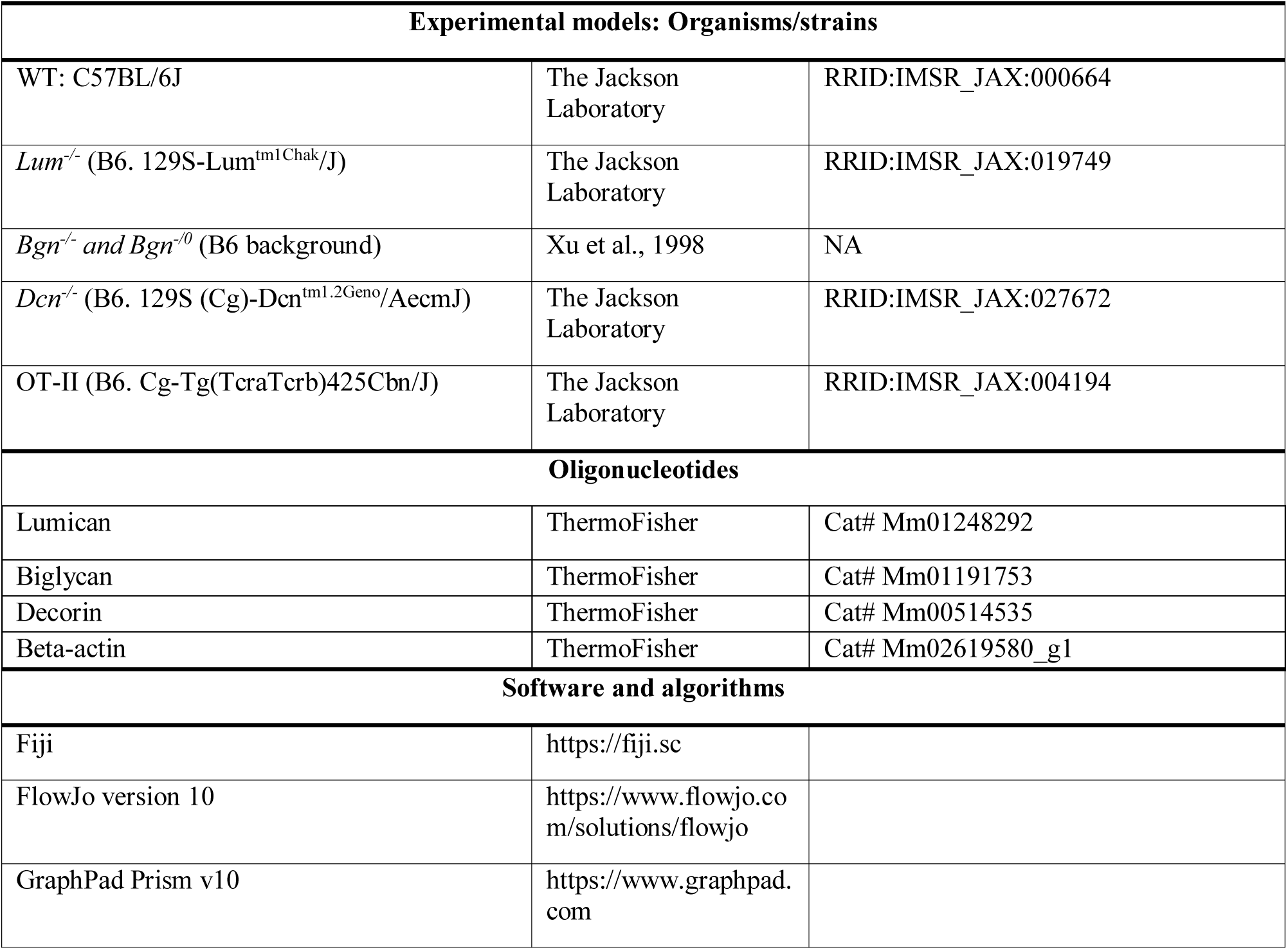
List of antibodies, reagents and resources.

