## Supplementary Materials for "Paracrine regulations of IFN-γ secreting CD4^+^ T cells by lumican and biglycan are protective in allergic contact dermatitis"

**This PDF file includes:**

Figs. S1 to S10

Table S1

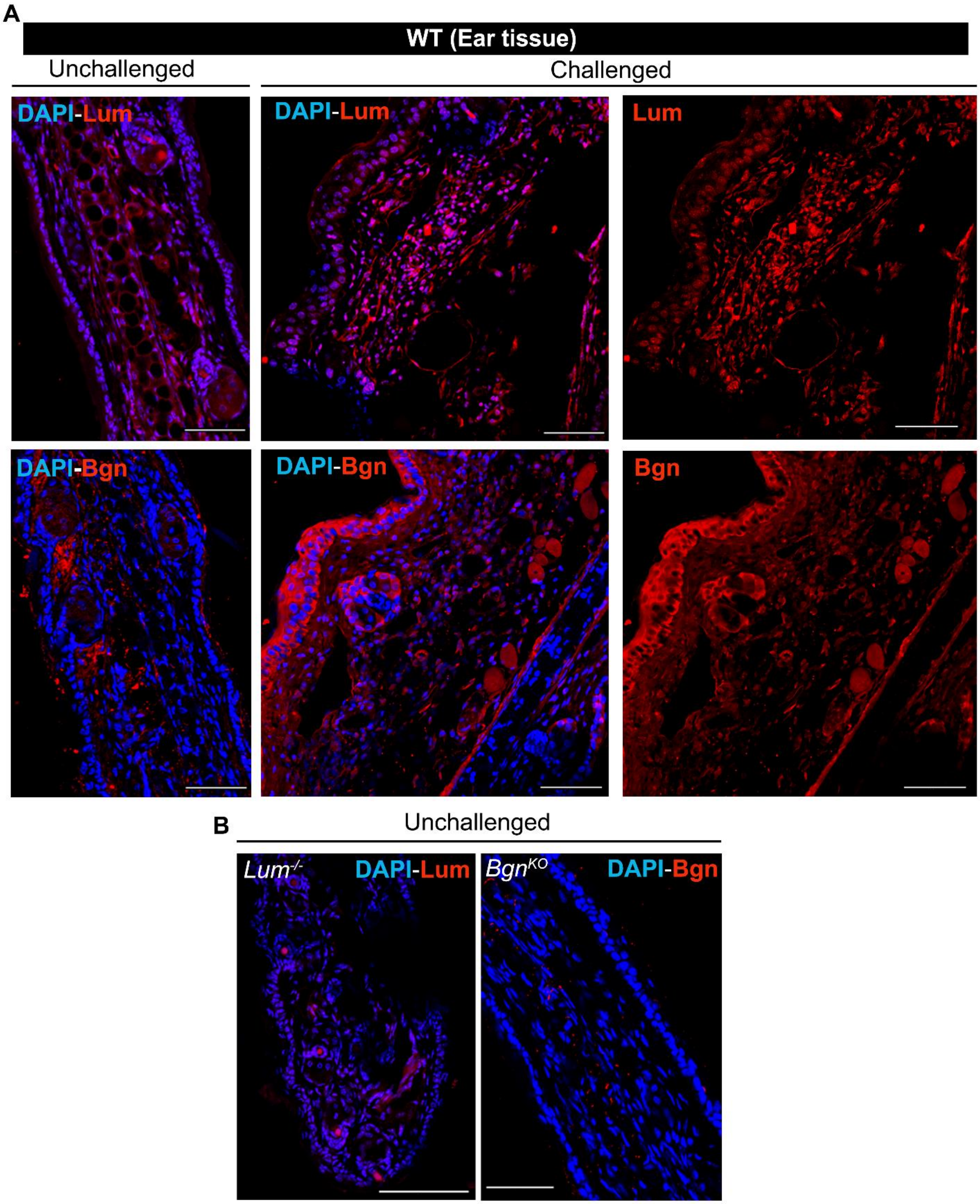


**Fig. S1: Lumican and biglycan expression in the WT ear tissue. (A)** Lumican (Lum) and biglycan (Bgn) level increases in the challenged WT ear tissue as shown by the IHC images. DAPI staining indicates the nucleus. **(B)** Ear tissue sections of *Lum*$^{-/-}$ and *Bgn*$^{KO}$ mouse served as a negative control for the primary antibodies and showed little to no immunoreactivity. Scale bars, 100 µm (A and B).

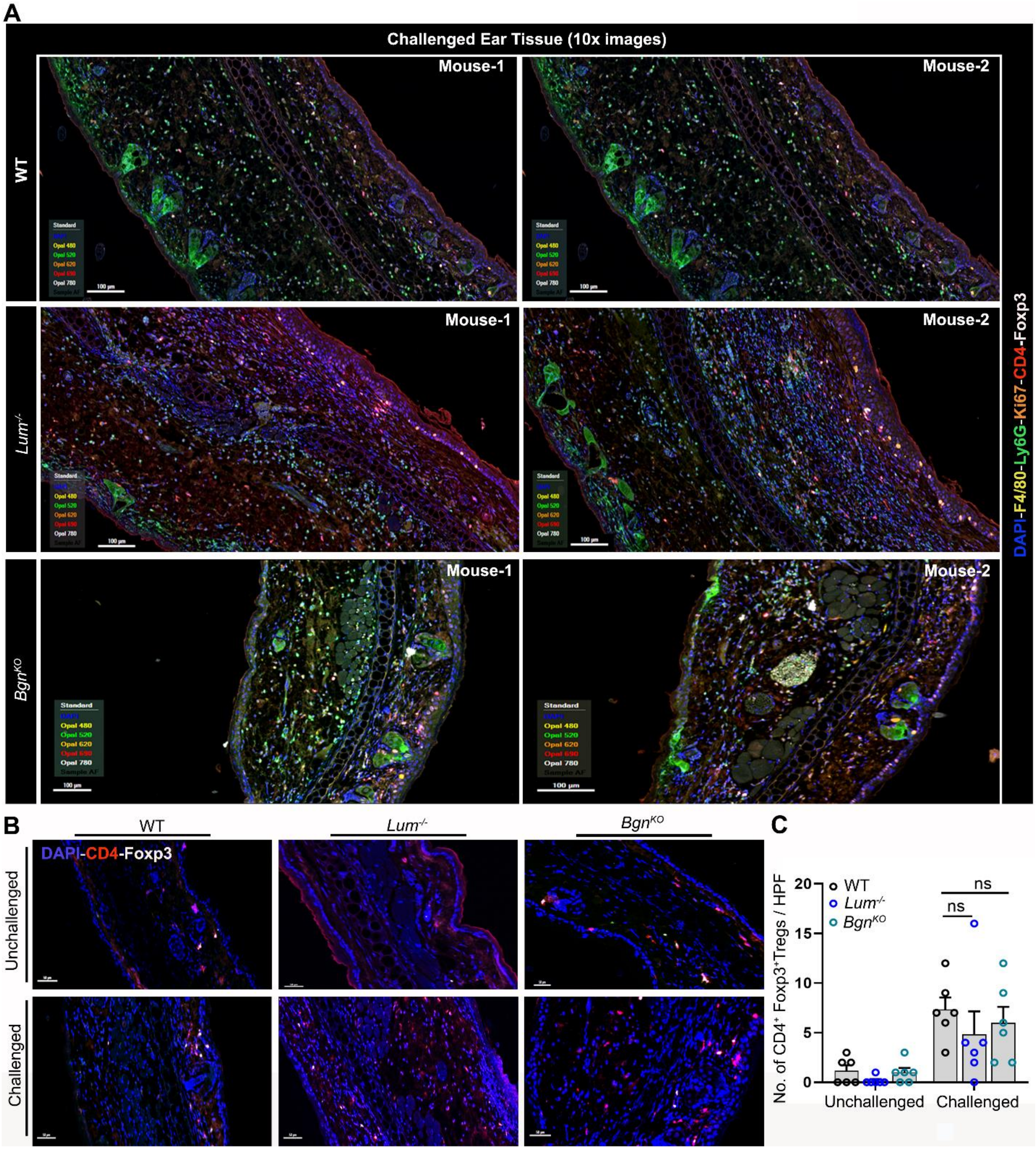


**Fig. S2: WT, *Lum*$^{-/-}$, *Bgn*$^{KO}$ have comparable number of Foxp3$^+$ Tregs in the ear tissue. (A-B)** Wild type and the null mice have similar number of CD4$^+$Foxp3$^+$ Tregs in the ear tissue as shown by the IHC images (A) and the cumulative bar graph from six high-power field (HPF) from one experiment with two mice per genotype (B). **(C)** IHC images showing all the different markers in the challenged ear tissue at 10x magnification from two mice per genotype. Scale bars, 100 µm (A and C). The error bars represent mean ± SEM, unpaired t test with Welch correction (B). *ns, not significant.*

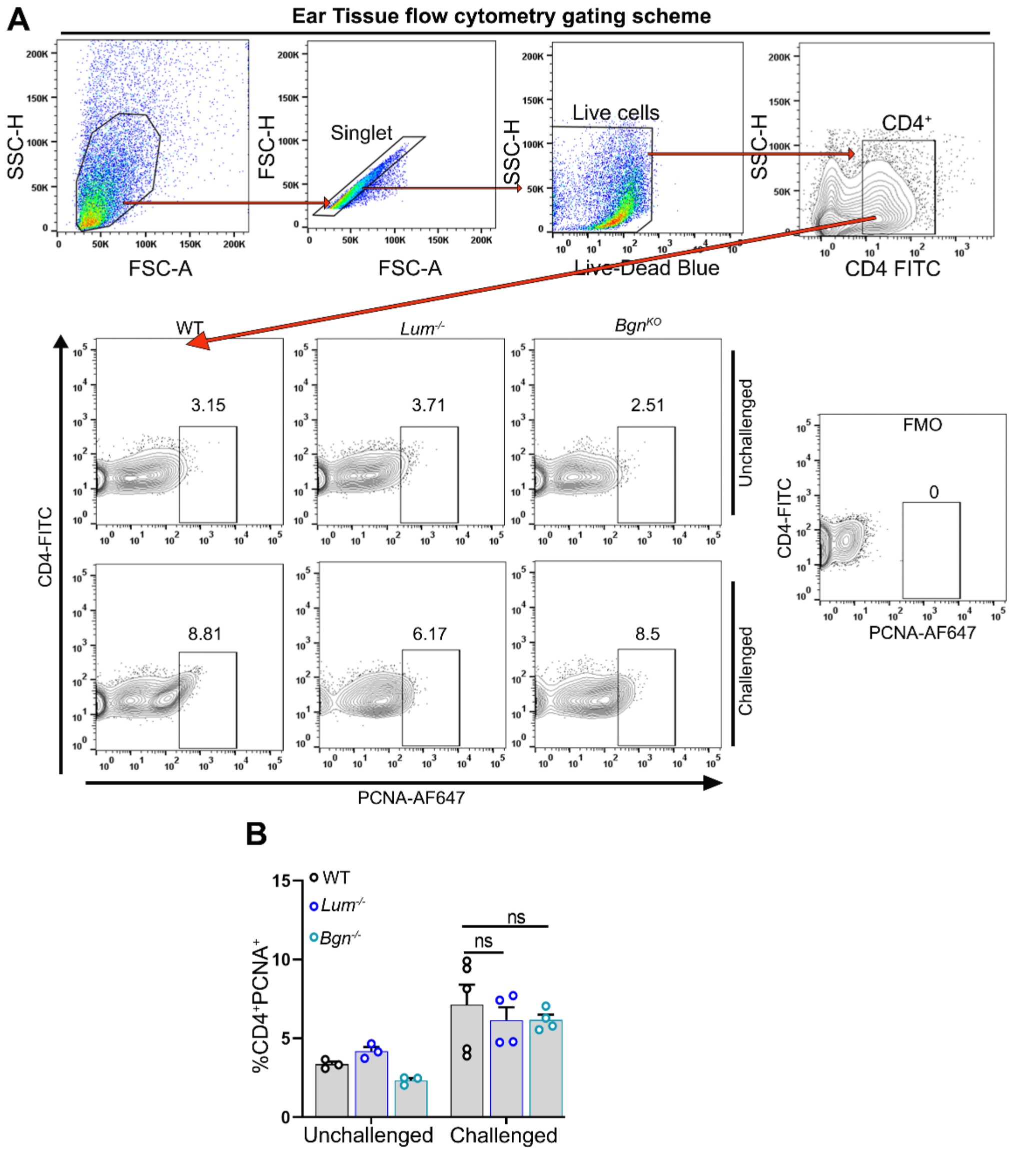


**Fig. S3: WT, *Lum*$^{-/-}$, *Bgn*$^{KO}$ have comparable percentage of proliferating CD4$^{+}$T cells in the ear tissue.** **(A)** Flow cytometry gating scheme for the proliferating PCNA$^{+}$CD4$^{+}$ T cells in the ear pinnae. **(B)** Cumulative data from 2 independent experiments (n = 3 – 5 mice/genotypes) shows the percentage of proliferating CD4$^{+}$ T cells (PCNA$^{+}$CD4$^{+}$) in the ear pinnae. The error bars represent mean ± SEM, two-tailed Student's t test. *ns, not significant*.

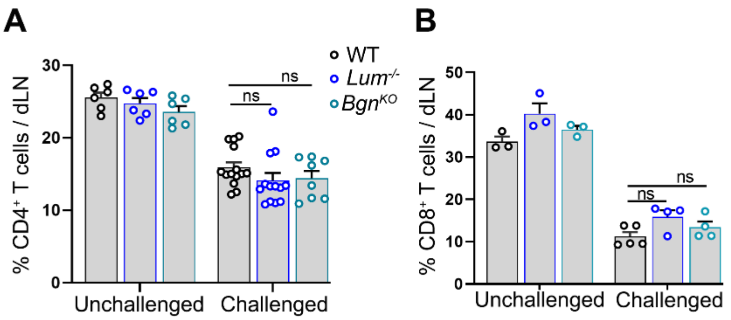


**Fig. S4: Percentage of $CD4^+$ and $CD8^+$ T cells in the dLN. (A)** The cumulative data shows the percentage of $CD4^+$ T cells (n = 6 – 14 mice from 3 independent experiments. **(B)** The cumulative data shows percentage of $CD8^+$ T cells in the dLN of unchallenged and challenged WT, *Lum*$^{-/-}$ and *Bgn*$^{KO}$ (n = 3 -5 mice from 2 independent). The error bars represent mean ± SEM, unpaired t test with Welch correction. *ns, not significant*.

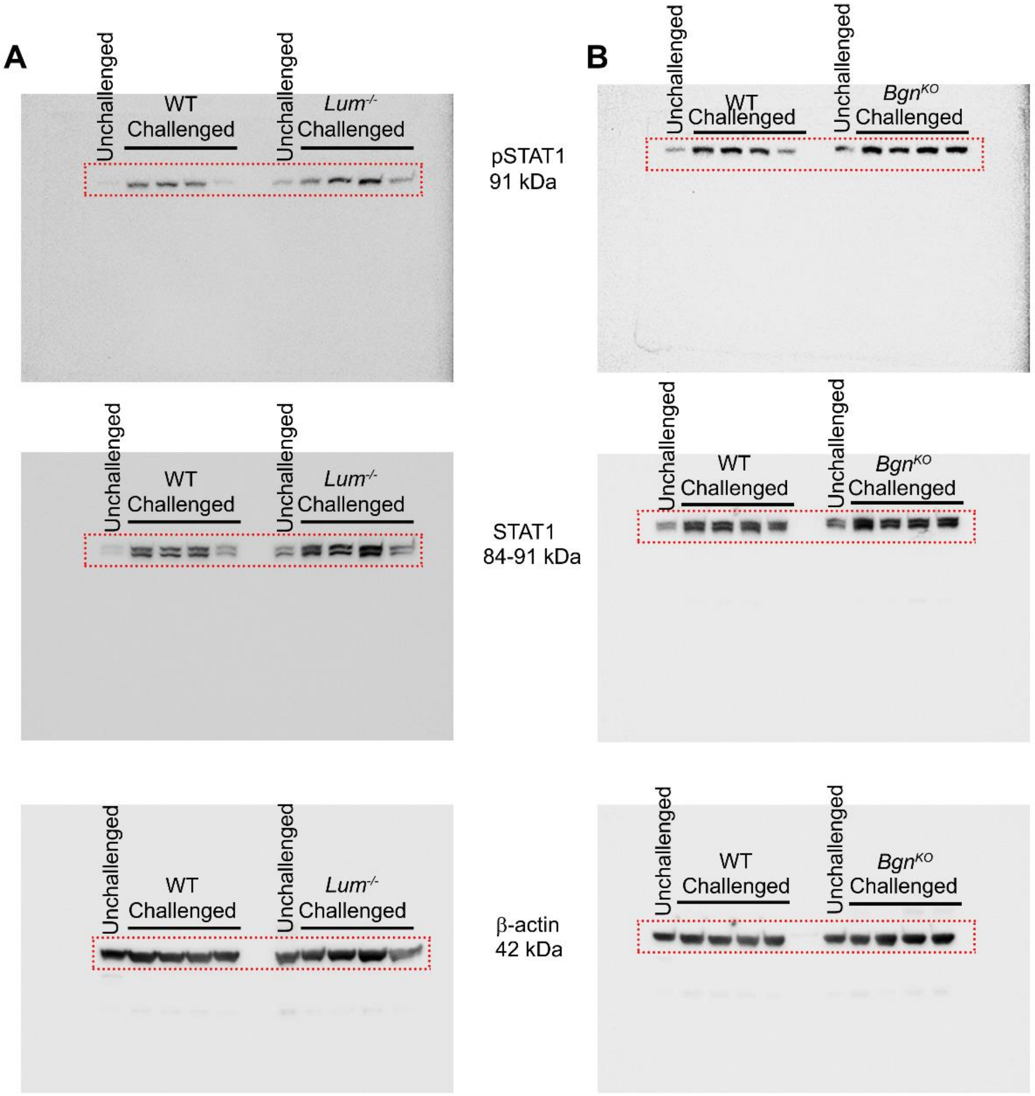


**Fig. S5: Full length Immunoblots showing phospho-STAT1, STAT1 and β-actin level in the dLN. (A and B)** Uncropped images of the blots probed for indicated proteins as presented in (A) for main Fig.5D and (B) for main Fig.5E.

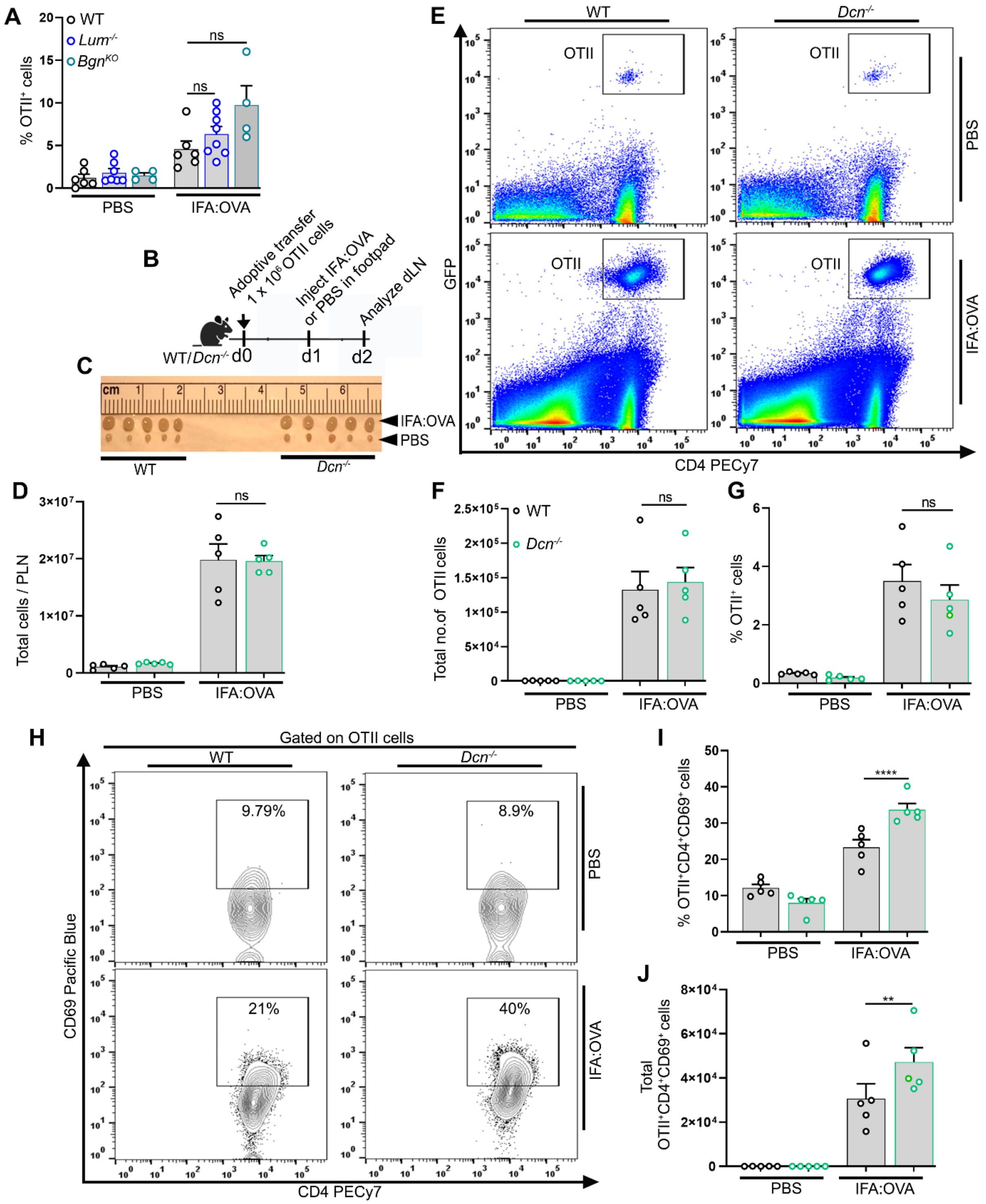


**Fig. S6: Decorin suppresses CD4$^+$ T cell activation *in vivo*. (A)** Cumulative data show percentage of transferred OT-II cells in the dLN (n = 4 – 8 mice/genotype) from two independent experiments. **(B)** Schematic showing the OT-II adoptive transfer model. On day0, GFP expressing OT-II cells (1 x 10$^6$) were adoptive transferred retro-orbitally to WT and *Dcn*$^{-/-}$ mice. Foot-pad were challenged with IFA-OVA or PBS on day1 and dLN were harvested for analysis on day2 after euthanasia. **(C and D)** Images showing similar sizes and comparable number of total cells in dLN of WT and *Dcn*$^{-/-}$ mice. **(E)** Representative dot plots showing transferred OT-II cells ($GFP^{+}CD4^{+}$ cells) in the dLN after injected with either IFA-OVA or PBS in WT and *Dcn*$^{-/-}$ mice. **(F and G)** Cumulative data show total number (E) and percentage (F) of transferred OT-II cells in the dLN (n = 5 mice/genotype) from one independent experiment. **(H)** Representative contour plots showing percentage of activated OT-II cells ($CD69^{+}CD4^{+}$ cells) in the dLN. **(I and J)** Cumulative data show the percentage (H) and total number (I) of activated OT-II cells ($CD69^{+}$OT-II$^{+}CD4^{+}$ cells) in the dLN after injected with either IFA-OVA or PBS in WT and *Dcn*$^{-/-}$ mice (n = 5 mice/genotype) from one experiment. The error bars represent mean ± SEM, unpaired t test with Welch correction. ***P < 0.01, ****P < 0.0001*; *ns, not significant*.

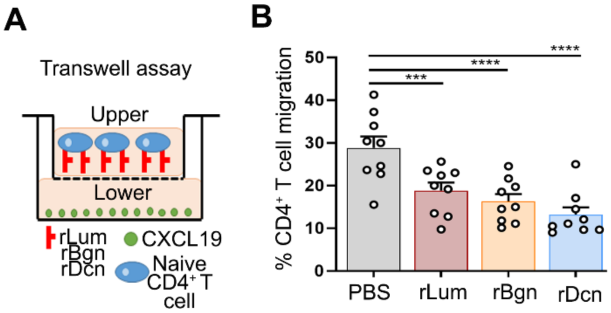


**Fig. S7: Lumican, biglycan and decorin suppresses CD4$^+$ T cell chemotaxis *in vitro*. (A)** Schematic shows the transwell migration assay. **(B)** Cumulative data from 2 independent experiments shows the number of CD4$^+$ T cells treated with 1 µg/ml of either rLum or rBgn or rDcn migrated towards 0.5 µg/ml rCCL19 in the lower chamber as percentage with respect to just media (no rCCL19). The error bars represent mean ± SEM, two-way ANOVA. ****P* < 0.001, *****P* < 0.0001.*

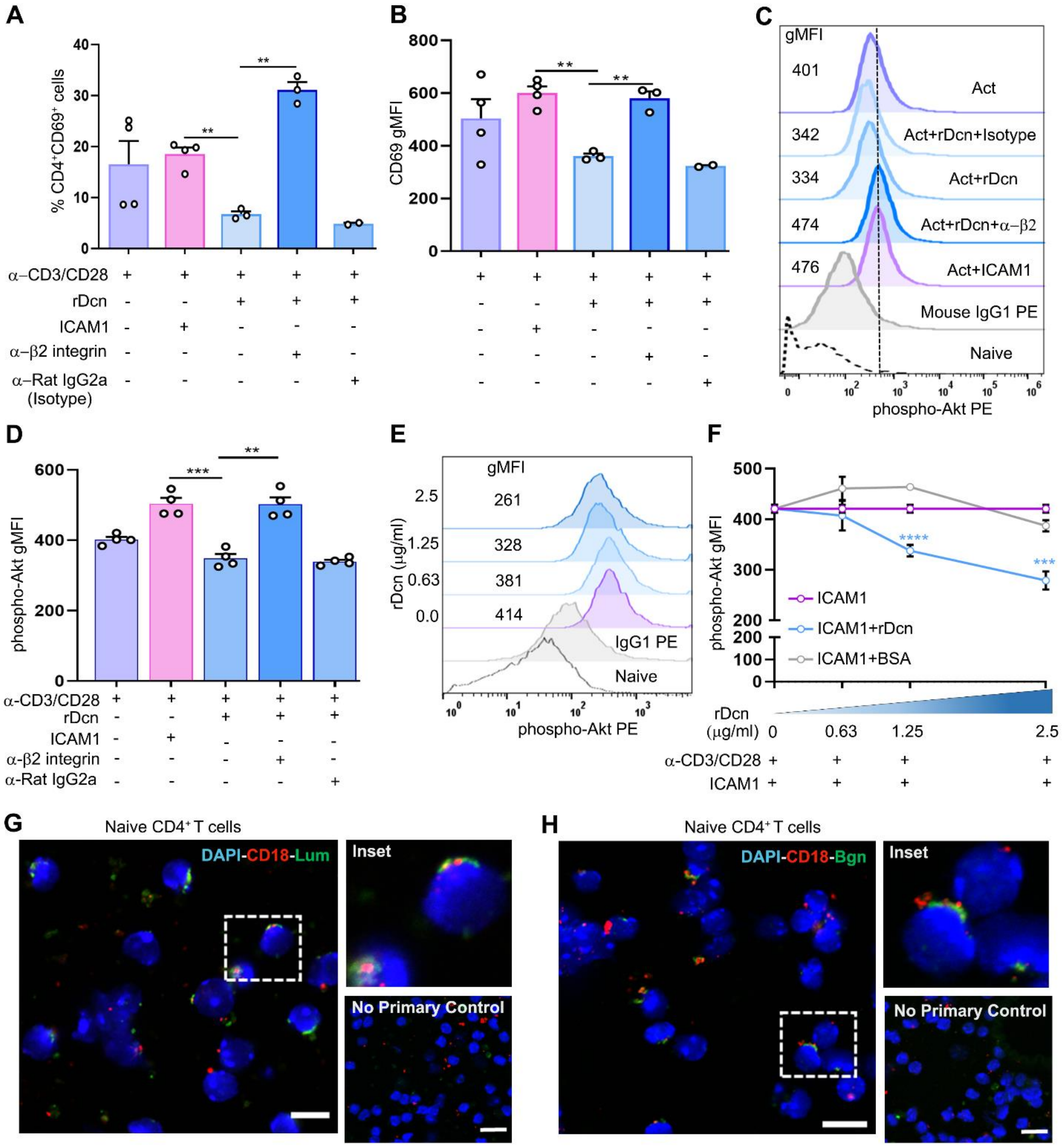

**Fig. S8: Decorin suppresses CD4$^+$ T cell activation. (A and B)** Cumulative data from 2 independent experiments shows the percentage (A) of activated CD4$^+$ T (CD69$^+$CD4$^+$) cells and surface expression of CD69 (gMFI) (B). **(C)** Representative histograms show the phospho-Akt-1 level in activated CD4$^+$ T cells with various treatments as indicated in the plot. **(D)** Cumulative data from 2 independent experiments shows the phospho-Akt-1 levels (gMFI) in activated CD4$^+$ T cells with either rDcn or ICAM-1 alone or in combination with anti-β2 integrin antibody. Anti-Rat IgG2a is used as an isotype control. **(E)** Representative histograms show the phospho-Akt-1 level in activated CD4$^+$ T cells as an indicative of ICAM-1 binding strength to LFA-1 in the presence of increasing soluble rDcn or BSA. **(F)** Cumulative line graph from 2 independent experiments shows the phospho-Akt-1 levels (gMFI) in activated CD4$^+$ T cells as a measure of competitive binding between LFA-1 and rDcn or ICAM-1. (**G** and **H**) Representative confocal image from 2 independent experiments shows staining of CD18 with rLum (G) and rBgn (H) in naïve $CD4^+$ T cells. No primary control indicates the specificity of primary antibodies. The histogram depicts cross-line scans of the fluorescence intensities of the merged panels. Scale bars, 10 µm. Each dot represents technical replicates (A, B and D). The error bars represent mean ± SEM, unpaired t test with Welch correction. ***P < 0.01, ***P < 0.0001, ****P < 0.0001*; *ns, not significant.*

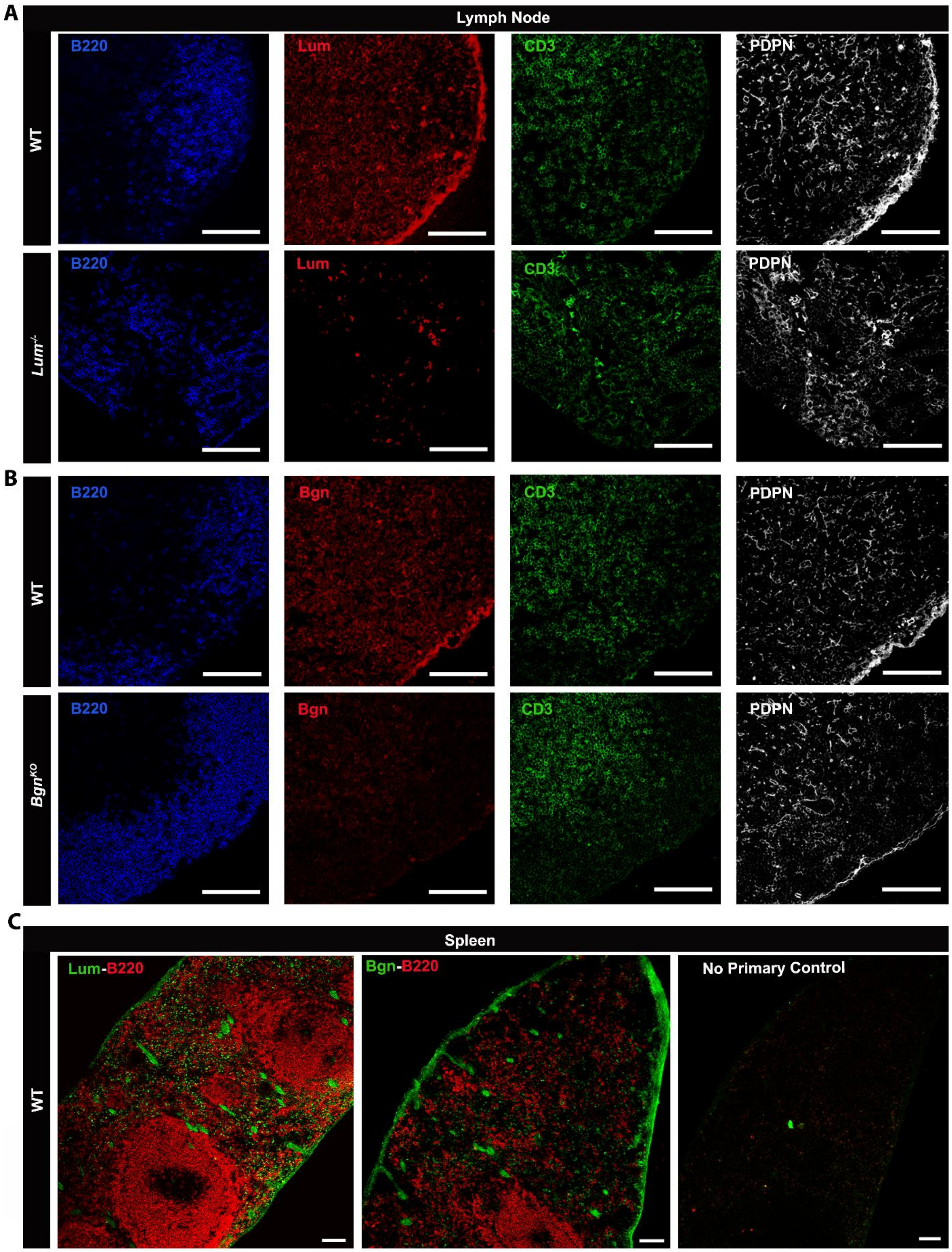


**Fig. S9: Localization of lumican and biglycan in the LN and spleen. (A and B)** Confocal images of showing individual channels for the merge images in main Figure 1B and 1C. **(C)**

Representative confocal images showing localization of lumican and biglycan, and B220 in the spleen of WT mice. Scale bars, 100 µm (A, B and C).

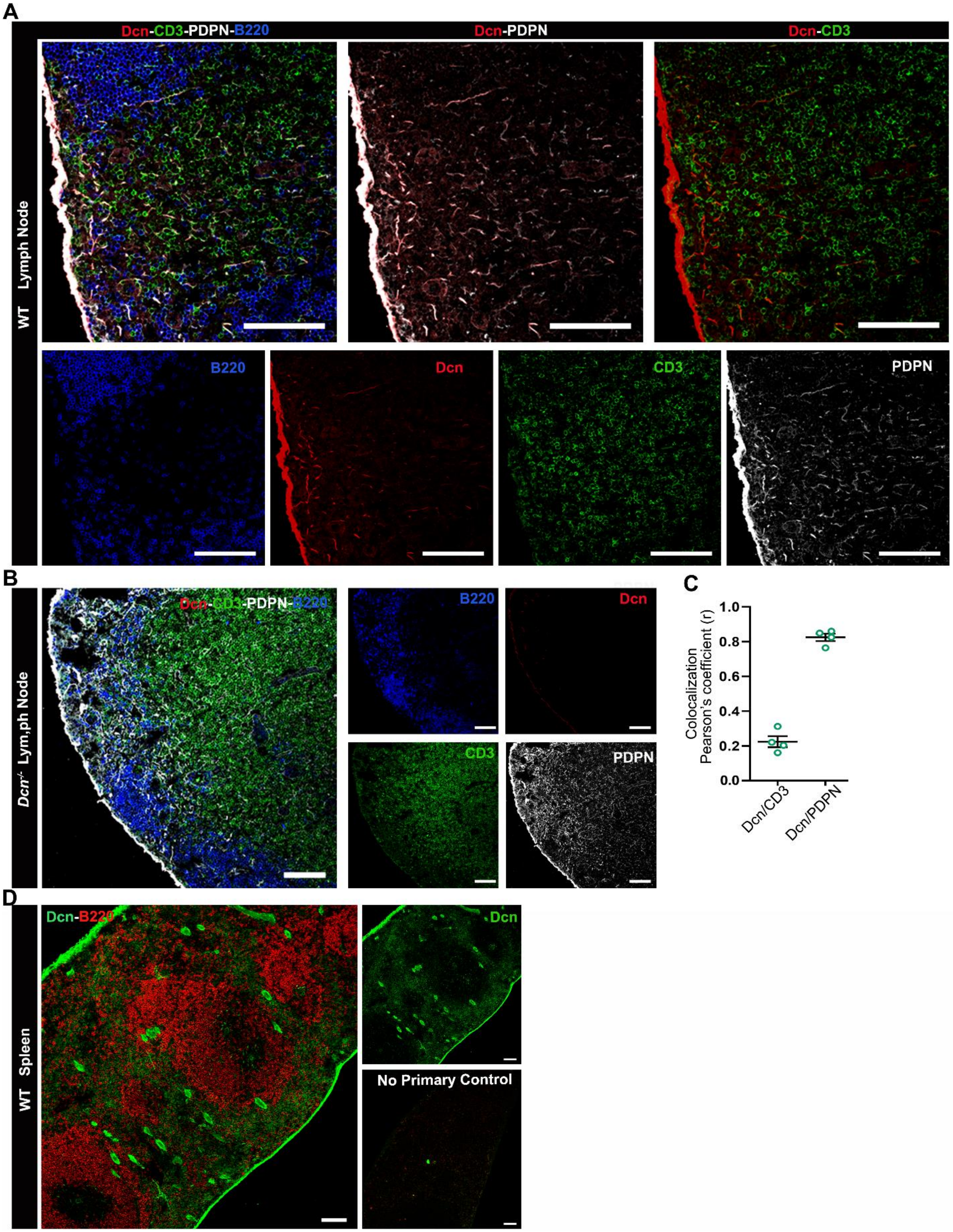


**Fig. S10: Localization of decorin in the LN and spleen. (A)** Representative confocal images showing localization of decorin with PDPN, B220 and CD3 in the lymph node regions of WT mice. **(B)** Representative confocal images showing the staining specificity of anti-mouse decorin antibody in the lymph node regions of *Dcn*$^{-/-}$ mice. **(C)** The cumulative data (n = 4 images) showing colocalization quantification of images in (A) by Pearson's correlation from 2 independent experiments. **(D)** Representative confocal images showing localization of decorin and B220 in the spleen of WT mice. Scale bars, 100 µm. The error bars represent mean ± SEM.

**Table S1: List of antibodies, reagents and resources.**

| REAGENT or RESOURCE | SOURCE | IDENTIFIER |
|---|---|---|
| **Antibodies** | | |
| Anti-mouse Lumican | Shao et al, 2012 | NA |
| Anti-mouse Lumican | R&D Systems | Cat # AF2745; RRID:AB_2139496 |
| Anti-mouse Biglycan | Larry Fisher/Kerafast | Cat # LF-159; RRID:AB_2920701 |
| Anti-mouse Decorin | Larry Fisher/Kerafast | Cat # LF-114; RRID:AB_3099478 |
| Anti-mouse PDPN, clone 8.1.1 | BioLegend | Cat # 127401; RRID:AB_1089187 |
| Anti-mouse B220, clone RA3-6B2 | eBioscience | Cat # 14-0452-82; RRID:AB_467254 |
| Anti-mouse CD3, clone 145-2C11 | eBioscience | Cat # 14-0031-82; RRID:AB_467049 |
| Anti-mouse CD3 APC, 17A2 | BioLegend | Cat # 100236; RRID:AB_2561455 |
| Anti-mouse CD4 FITC, clone GK1.5 | BioLegend | Cat # 100406; RRID:AB_312690 |
| Anti-mouse CD4 PECy7, clone GK1.5 | BioLegend | Cat # 100421; RRID:AB_312706 |
| Anti-mouse CD4 BV421, clone GK1.5 | BioLegend | Cat # 100437; RRID:AB_2562557 |
| Anti-mouse CD8 PECy7, clone 53-6.7 | BioLegend | Cat # 100721; RRID:AB_312760 |
| Anti-mouse CD8 PE, clone 53-6.7 | BioLegend | Cat # 100708; RRID:AB_312747 |
| Anti-mouse CD69 APC, clone H1.2F3 | BioLegend | Cat # 104513; RRID:AB_492843 |
| Anti-mouse CD69 Pacific Blue, clone H1.2F3 | BioLegend | Cat # 104524; RRID:AB_2260064 |
| Anti-mouse PCNA AF647, clone PC10 | BioLegend | Cat # 307912; RRID:AB_2267947 |
| Anti-mouse Ly6G PE, clone 1A8 | BioLegend | Cat # 127607; RRID:AB_1186104 |
| Anti-mouse F4/80 FITC, clone BM8 | BioLegend | Cat # 123108; RRID:AB_893502 |
| Anti-mouse Ly6C BV605, clone HK1.4 | BioLegend | Cat # 128035; RRID:AB_2562352 |
| Anti-mouse CD11b APC, clone M1/70 | BioLegend | Cat # 101211; RRID:AB_312794 |
| Anti-mouse CD11c BV786, clone N418 | BioLegend | Cat # 117335; RRID:AB_11219204 |
| Anti-mouse IL-17 APC, clone TC11-18H10.1 | BioLegend | Cat # 506915; RRID:AB_536017 |
| Anti-mouse IFN-y BV711, clone XMG1.2 | BioLegend | Cat # 505835; RRID:AB_11219588 |
| Anti-mouse IL-4 PE, clone 11B11 | BioLegend | Cat # 504103; RRID:AB_315318 |
| Anti-mouse T-bet PE, clone 4B10 | BioLegend | Cat # 644809; RRID:AB_2028583 |

| | | |
|---|---|---|
| Anti-mouse phospho-ZAP-70 PE, clone J.947.4 | Invitrogen | Cat # MA5-28072; RRID:AB_2745063 |
| Anti-mouse phospho-Lck PE, clone A18002D | BioLegend | Cat # 933103; RRID:AB_2820204 |
| Anti-mouse phospho-Akt PE, clone A21001C | BioLegend | Cat # 606554; RRID:AB_3068251 |
| Anti-mouse CD25 PE, clone 3C7 | BioLegend | Cat # 101903; RRID:AB_312847 |
| Anti-mouse β2 integrin (CD18), clone M18/2 | BioLegend | Cat # 101417; RRID:AB_2832276 |
| Anti-Rat IgG2a, clone RTK2758 | BioLegend | Cat # 400543; RRID:AB_11148951 |
| Anti-mouse CD28, clone 37.51 | BioLegend | Cat # 102112; RRID:AB_312877 |
| Anti-mouse CD3, clone 145-2C11 | BioLegend | Cat # 100340; RRID:AB_2616674 |
| Anti-mouse STAT-1, clone D1K9Y | Cell Signaling | Cat # 14994; RRID:AB_2737027 |
| Anti-mouse phospho-STAT-1 | Cell Signaling | Cat # 9177; RRID:AB_2197983 |
| Anti-mouse beta-actin, clone 8H10D10 | Cell Signaling | Cat # 3700; RRID:AB_2242334 |
| Anti-mouse Foxp3 AF488, clone MF-14 | BioLegend | Cat # 126406; RRID:AB_1089113 |
| Anti-mouse Ki67 APC, clone 16A8 | BioLegend | Cat # 652409; RRID:AB_2562141 |
| Anti-Syrian Hamster-AF 647 | Abcam | Cat # ab180117; RRID:AB_3099479 |
| Anti-Armenian Hamster-AF 488 | Abcam | Cat # ab173003; RRID:AB_2936402 |
| Anti-Goat-AF 555 | Abcam | Cat # ab150134; RRID:AB_2715537 |
| Anti-Rat-AF 405 | Abcam | Cat # ab175670; RRID:AB_3099480 |
| Anti-Rabbit-AF 555 | Abcam | Cat # ab150086; RRID:AB_2890032 |
| Anti-6X-His Tag Monoclonal antibody-AF 488 | Invitrogen | Cat # MA1-135-A488; RRID:AB_2610635 |
| Anti-Rat-NL 557 | R&D Systems | Cat # NL013; RRID:AB_884217 |
| Anti-Rabbit IgG - HRP | Cell Signaling | Cat # 7074; RRID:AB_2099233 |
| Anti-Mouse IgG - HRP | Cell Signaling | Cat # 7076; RRID:AB_330924 |
| **Chemicals, peptides, and recombinant proteins** | | |
| Protease Inhibitor Cocktail | Roche | Cat # 11836170001 |
| 2, 4-dinitrofluorobenzene (DNFB) | Sigma-Aldrich | Cat # D1529 |
| Acetone | Sigma-Aldrich | Cat # 67-64-1 |
| OVA peptide | Sigma-Aldrich | Cat # O1641 |

| | | |
|---|---|---|
| Incomplete Freund Adjuvant (IFA) | Sigma-Aldrich | Cat # AR002 |
| Liberase TL | Roche | Cat # 05401020001 |
| DNase 1 | Sigma-Aldrich | Cat # DN25 |
| beta-Mercaptoethanol (BME) | Sigma-Aldrich | Cat # 444203 |
| Trypan Blue Exclusion Dye | Sigma-Aldrich | Cat # 15250061 |
| Phorbol 12-myristate 13-acetate (PMA) | Sigma-Aldrich | Cat # P1585 |
| Ionomycin | Sigma-Aldrich | Cat # I0634 |
| Brefeldin-A | BioLegend | Cat # 420601 |
| Dispase II | Sigma | Cat # D4693 |
| Collagenase P | Roche | Cat # 11249002001 |
| EasySep Buffer | Stemcell Technologies | Cat # 20144 |
| Mojosort Buffer 5X | Biolegend | Cat # 480017 |
| RPMI 1640 | Gibco | Cat # 11875093 |
| DMEM | Gibco | Cat # 11965092 |
| Alpha-MEM | Gibco | Cat # 12571063 |
| 100x Antibiotic/Antimycotic | Gibco | Cat # 15240062 |
| Fetal Bovine Serum | Gibco | Cat # 16140071 |
| Charcoal Stripped Serum | Gibco | Cat # A3382101 |
| 16% Paraformaldehyde (PFA) | ThermoFisher | Cat # 28908 |
| Recombinant human Lumican | In house | NA |
| Recombinant mouse Lumican | R&D Systems | Cat # 2745-LU |
| Recombinant mouse Biglycan | R&D Systems | Cat # 8128-CM |
| Recombinant mouse Decorin | R&D Systems | Cat # 1060-DE |
| Recombinant mouse ICAM-1 | R&D Systems | Cat # 796-IC |
| Recombinant mouse CCL19 | R&D Systems | Cat # 440-M3 |
| DAPI | BD Biosciences | Cat # 564907 |
| DTT | Sigma-Aldrich | Cat # D0632 |
| Vectashield antifade mounting media | Vector Laboratories | Cat # H-1000 |
| Normal Goat Serum | Sigma-Aldrich | Cat # D9663 |
| Normal Donkey Serum | Sigma-Aldrich | Cat # G9023 |

| | | |
|---|---|---|
| Taqman Fast advance Mastermix | ThermoFisher | Cat # 4444963 |
| NuPAGE 4-12% Bis-Tris gel | Invitrogen | Cat # NP0322BOX |
| NuPAGE MOPS SDS running buffer | Invitrogen | Cat # NP0001 |
| Intracellular Staining Permeabilization Wash Buffer | BioLegend | Cat # 421002 |
| True-Phos Perm Buffer | BioLegend | Cat # 425401 |
| True-Nuclear Transcription Factor Buffer Set | BioLegend | Cat # 424401 |
| **Critical commercial assays** | | |
| EasySep Mouse CD4+ T cell Isolation Kit | StemCell Technologies | Cat # 19852 |
| MojoSort Mouse CD4 Naïve T cell isolation Kit | BioLegend | Cat # 480039 |
| MojoSort Mouse CD45 Nanobeads | BioLegend | Cat # 480027 |
| Live/Dead Fixable Blue dead cell stain kit | ThermoFisher | Cat # L23105 |
| RNAeasy Mini Kit | Qiagen | Cat # 74104 |
| cDNA Synthesis kit | Bio-Rad | Cat # 1708891 |
| ArC Amine Reactive Compensation Bead Kit | ThermoFisher | Cat # A10628 |
| UltraComp eBeads Plus Compensation Beads | ThermoFisher | Cat # 01-3333-42 |
| **Deposited data** | | |
| None | | |
| **Experimental models: Organisms/strains** | | |
| WT: C57BL/6J | The Jackson Laboratory | RRID:IMSR_JAX:000664 |
| *Lum*$^{-/-}$ (B6. 129S-Lum$^{tm1Chak}$/J) | The Jackson Laboratory | RRID:IMSR_JAX:019749 |
| *Bgn*$^{-/-}$ *and Bgn*$^{-/0}$ (B6 background) | Xu et al., 1998 | NA |
| *Dcn*$^{-/-}$ (B6. 129S (Cg)-Dcn$^{tm1.2Geno}$/AecmJ) | The Jackson Laboratory | RRID:IMSR_JAX:027672 |
| OT-II (B6. Cg-Tg(TcraTcrb)425Cbn/J) | The Jackson Laboratory | RRID:IMSR_JAX:004194 |
| **Oligonucleotides** | | |

| | | |
|---|---|---|
| Lumican | ThermoFisher | Cat# Mm01248292 |
| Biglycan | ThermoFisher | Cat# Mm01191753 |
| Decorin | ThermoFisher | Cat# Mm00514535 |
| Beta-actin | ThermoFisher | Cat# Mm02619580_g1 |
| **Software and algorithms** | | |
| Fiji | https://fiji.sc | |
| FlowJo version 10 | https://www.flowjo.com/solutions/flowjo | |
| GraphPad Prism v10 | https://www.graphpad.com | |
